# A proper Excitatory/Inhibitory ratio is required to develop synchronized network activity in mouse cortical cultures

**DOI:** 10.1101/2025.02.28.640720

**Authors:** Eleonora Crocco, Ludovico Iannello, Fabrizio Tonelli, Gabriele Lagani, Luca Pandolfini, Giuseppe Amato, Angelo Di Garbo, Federico Cremisi

## Abstract

Excitatory/inhibitory (E/I) balance is thought to play a key role in cortical activity development. However, the modeling of cortical networks with different E/I ratios is not feasible *in vivo*. To address this point, we modeled an *in vitro* cortical network deployed of the inhibitory neurons normally migrating from the ventral telencephalon. Moreover, we implemented striatal cultures and co-cultures with mixed proportions of cortical and striatal neurons. The resulting cultures contained various proportions of inhibitory Parvalbumin (PV)^+^ neurons, ranging from 7% to 73%. Interestingly, these pure and mixed cortical/striatal cultures exhibited four distinct patterns of spontaneous activity and functional connectivity. Our findings highlighted a critical role for the inhibitory component in developing correlated network activity. Unexpectedly, cortical networks with 7% of PV^+^ neurons were not able to generate appreciable network burst activity due to the development of a strong network inhibition, despite their lowest E/I ratio. Our observations support the notion that an optimal ratio of PV^+^ neurons during cortical development is essential for the establishment of local inhibitory networks capable of generating and spreading correlated activity.

**Highlights:**

- *In vitro* neurogenesis models the development of mouse cortical network devoid of inhibitory neurons
- Cortical network with low inhibitory neuron ratio develops poor synchronized network activity
- GABA inhibition unmask intrinsic network ability to generate highly synchronized activity
- Network response to single node stimulus depends on optimal inhibitory neuron ratio
- A proper excitatory/inhibitory ratio is necessary for the development of network burst activity

**Graphical Abstract:** 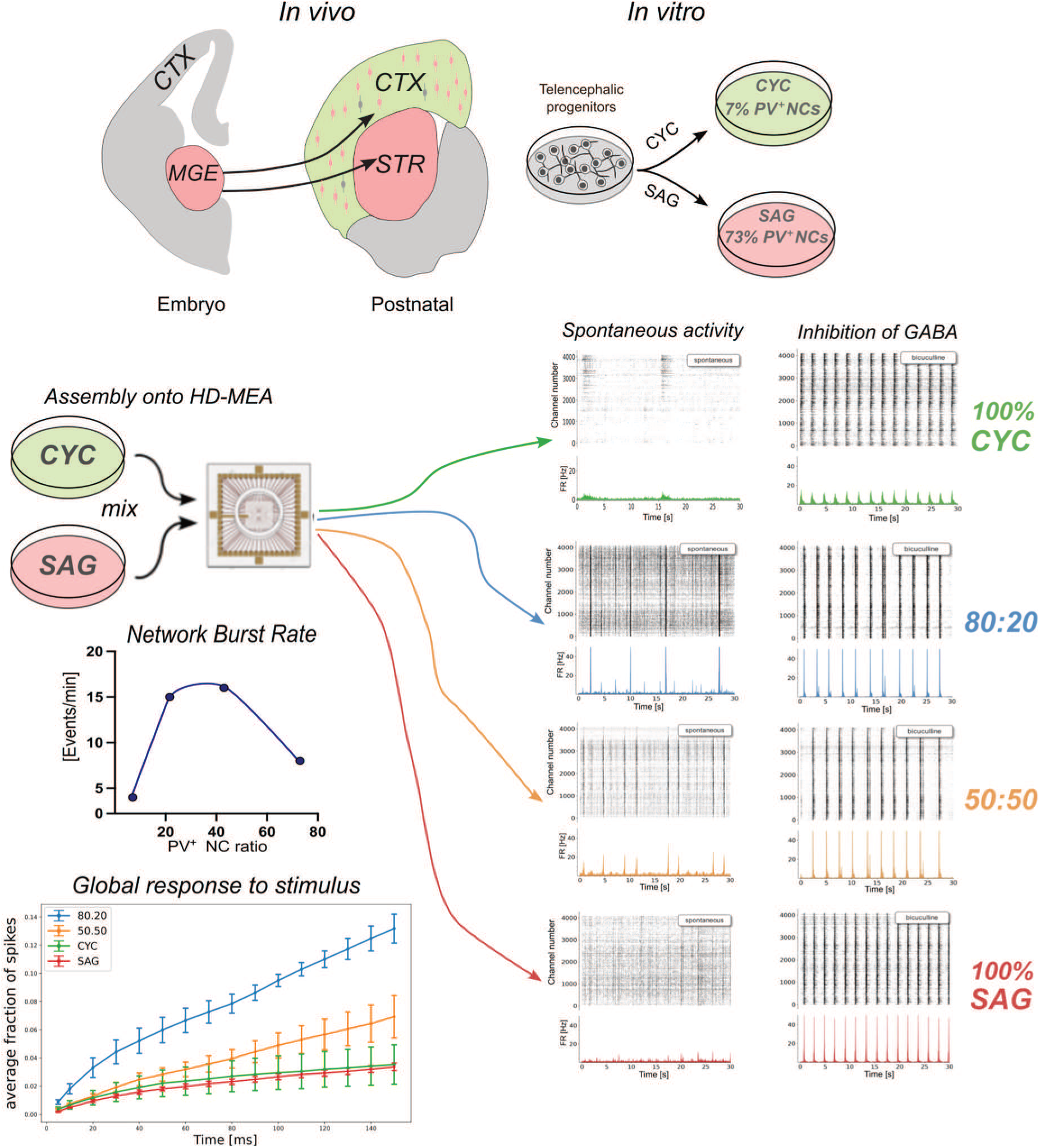

## INTRODUCTION

The cerebral cortex contains dozens of different types of neurons that populate specific cortical areas and layers and are generally divided into two main classes: glutamatergic pyramidal neurons, which constitute on average 85% of cortical neurons and send long-range projections to other cortical or subcortical targets, and GABAergic interneurons, which account for the remaining 15% of cortical neurons and exhibit only local connectivity (Molyneaux et al., 2007). Pyramidal neurons and interneurons can be further subdivided into subtypes with specific functions and molecular properties (Lim et al., 2018; Van Den Ameele et al., 2014).

Mouse embryonic stem cells (mESCs) or human induced pluripotent cells (hiPSCs) have been used to model the early development of distinct encephalic regions *in vitro* by controlling different signaling pathways in precise time windows during their neuralization (Chambers et al., 2009; Chiaradia and Lancaster, 2020). This approach enables the generation of both glutamatergic and GABAergic cortical neurons and, in principle, the generation of cultured neuronal networks with different proportions of the two types.

Forebrain identity is thought to constitute a primitive pattern of neural identity, which is acquired and retained through local inhibition of caudalizing morphogen signals, primarily BMP and Wnt (Bertacchi et al., 2015; Watanabe et al., 2005a; Ying et al., 2003). Dorsalization of telencephalic progenitors can be achieved *in vitro* using the Sonic Hedgehog (Shh) inhibitor, Cyclopamine, which induces differentiation into cortical pyramidal neurons (Gaspard et al., 2008). On the other hand, specification of ventral telencephalic cells requires Shh activation (Li et al., 2009; Watanabe et al., 2005b), which can be obtained *in vitro* via administration of the Shh agonist, SAG (Cederquist et al., 2019; Danjo et al., 2011), leading to the production of ventral telencephalic interneuron progenitors (Xu et al., 2010). The mature cerebral cortex is established only after the migration of these inhibitory neuronal progenitors from ventral telencephalic regions (mainly from the medial ganglionic eminence, MGE) into the dorsal telencephalic aspect (Wonders and Anderson, 2006). After migration, the GABAergic inhibitory interneurons integrate and form synaptic connections with glutamatergic excitatory neurons, establishing a balanced Excitation/Inhibition (E/I) ratio, which is crucial for the normal brain function (Gelman and Marín, 2010; Kepecs and Fishell, 2014; Lodato et al., 2011). Conversely, an unbalanced E/I ratio in the cortex may lead to altered circuitry and reduced information processing, and has been implicated in several brain disorders such as Schizophrenia or Autism Spectrum Disorders (ASDs) (Lee et al., 2017; Nelson and Valakh, 2015; Sohal and Rubenstein, 2019).

To investigate neural networks with different E/I ratios and functionally characterize them during developmental maturation, we modeled two distinct populations of cells *in vitro*, namely dorsal telencephalic and striatal progenitors. This approach enabled us to examine how cortical network activity developed in the absence of ventral inhibitory neurons. We compared the activity of a network composed of dorsal telencephalic neurons with that of a striatal network and with networks formed by varying ratios of dorsal and ventral telencephalic neurons. Finally, we analyzed basic structural and functional network parameters such as synapse density, firing activity, network burst synchronization and connectivity, together with the ability to respond to electrical stimulations. Our findings indicate that the E/I balance dramatically affects the development and maturation of neuronal cultures, highlighting crucial differences of network connectivity and activity of cultures. Overall, we observed that a proper E/I ratio is required for the formation and spreading of correlated activity in cortical networks.

## RESULTS

### Cyclopamine and SAG treatments respectively induce the production of cortical and striatal neurons

During the first week of mESC neuralization, early double inhibition of BMP and Wnt signaling (WiBi) allows the generation of a general dorsal telencephalic identity (Bertacchi et al., 2015; Terrigno et al., 2018). Based on this, we established a protocol to specifically derive dorsal and ventral telencephalic neurons from mESCs (Figure 1A,B). To induce a dorsal cortical identity, together with WiBi, from DIV5 to DIV10 we added Cyclopamine, a known antagonist of the Shh developmental pathway (Chen et al., 2002a, Gaspard et al. 2008). On the other hand, to obtain a ventral identity, during the same time window, we added SAG, a Shh agonist that acts directly on Smoothed receptor (Chen et al., 2002b; Frank-Kamenetsky et al., 2002). First, we assessed the positional identity of these cells at DIV11, an early stage of neural differentiation when markers of telencephalic and subpallial identity are specifically expressed in the developing embryo (Fuccillo et al., 2006; Kobayashi et al., 2002; Martynoga et al., 2005). We started by observing the expression of general telencephalic/positional markers, such as *Pax6, Nkx2.1, FoxG1* (Figure 1B-E, S1A-C). Cyclopamine-treated (CYC) cells expressed high levels of the dorsal marker *Pax6,* while SAG-treated (SAG) cells exhibited a high expression of the ventral identity marker *Nkx2.1*. Moreover, both types of cells had high levels of *FoxG1* expression compared to control neuralized cells without Wnt inhibition (Midbrain), confirming the establishment of a telencephalic identity.

**Figure 1.**
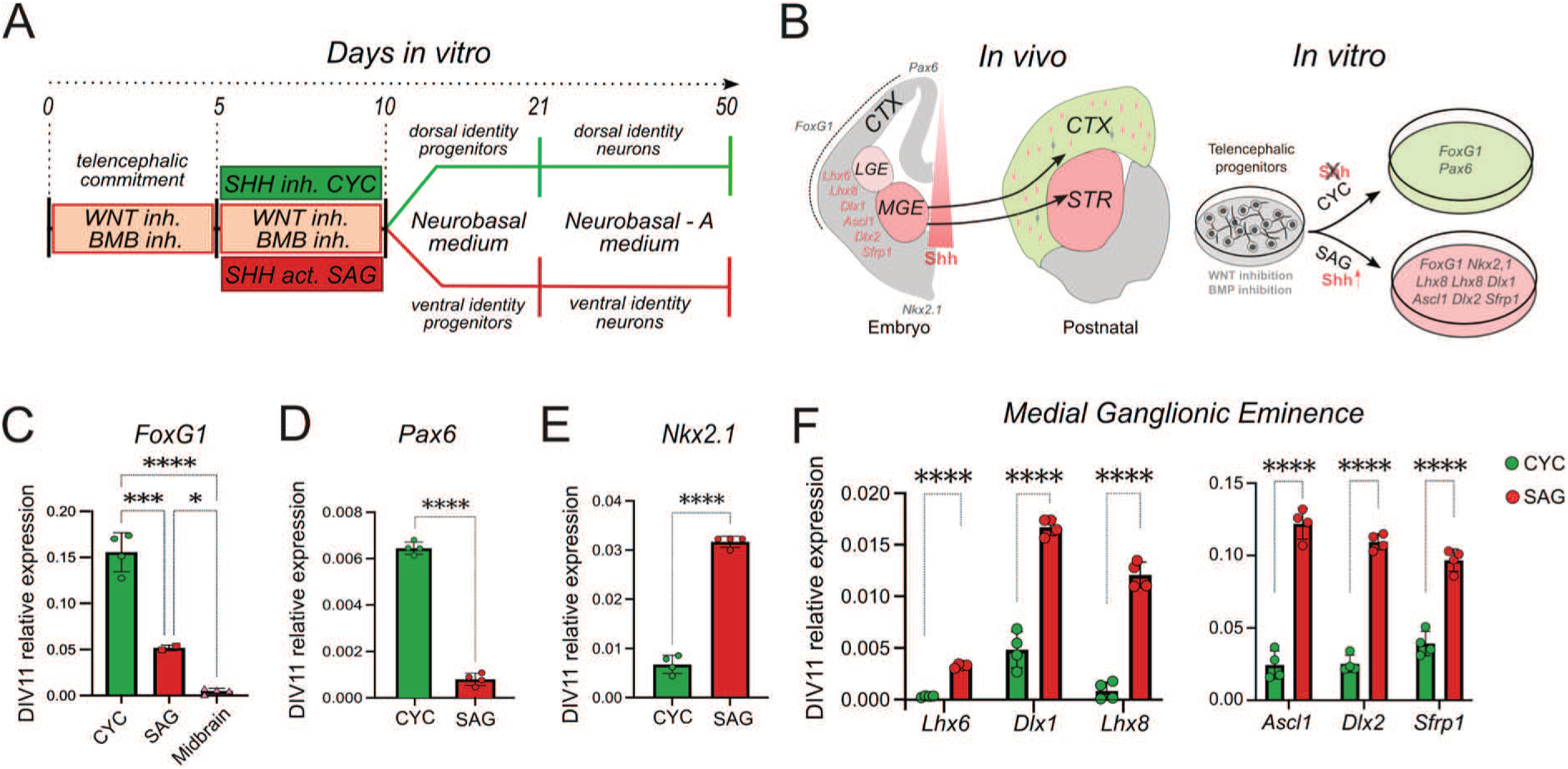
Positional identity and characterization of CYC and SAG progenitor cells. A) Protocol of mESC neuralization. Medium compositions are reported in Methods. B) Schematic representation of the expression of the genes analyzed in Figure 1B-E *in vivo* and their respective expression in CYC and SAG cultures *in vitro*. CTX: cortex, STR: striatum; MGE: medial ganglionic eminence; LGE: lateral ganglionic eminence. C-D-E) Relative expression of early pallial markers evaluated by qRT-PCR in the two different cultures (n = 4 independent experiments). Midbrain in C): cultures without WiBi-induced neuralization with mesencephalic identity (Bertacchi et al., 2015). Mean ± SD is shown; ordinary one-way ANOVA with Tukey’s multiple comparisons test was performed for *Foxg1* expression; unpaired t-test was performed for *Pax6* and *Nkx2.1* markers. F) Relative expression of MGE-LGE markers evaluated by qRT-PCR in the two different cultures (n = 4 independent experiments). Mean ± SD is shown; Multiple unpaired t-test with Holm-Šídák correction method. P-values: *p-value < 0.05, ***p-value < 0.001, ****p-value < 0.0001.

We proceeded to analyze specific early markers of the subpallium and medial and lateral ganglionic eminences (MGE-LGE), such as *Lhx6, Lhx8 and Dlx1* (Chen et al., 2017), and *Ascl1, Dlx2 and Sfrp1* (Nery et al., 2002). We found an increased expression of these genes in SAG cells compared to the other neural population (Figure 1F, S1D-I). These data confirmed that SAG cultures have a ventral identity similar to the MGE and subpallium. A more detailed analysis performed at later stages of differentiation highlighted the different nature of neurons produced by the two cultures. Since the MGE is the source of a wide variety of interneurons (Gelman et al., 2011; Xu et al., 2004), we evaluated specific subtype markers such as Parvalbumin, SST and VIP (Figure 2A). In particular, at DIV30 we noticed a significantly higher transcription of Parvalbumin (*Pvalb*) gene in SAG-treated cultures compared to CYC-treated cultures, in line with the prevalence of Parvalbumin-expressing neurons generated in MGE and migrated into the cortex (Lim et al., 2018; Wonders and Anderson, 2006). We confirmed the presence of Parvalbumin^+^ (PV) neurons by immunofluorescence analysis at DIV35 (Figure 2B). We counted 73% PV^+^ neurons in SAG-treated cultures, while we found a much lower percentage (7%) in CYC cultures. Moreover, given the known association of PV^+^ interneurons with Perineuronal Nets (PNNs) (Fawcett et al., 2019; Lupori et al., 2023), we investigated also the reactivity against Wisteria floribunda agglutinin (WFA), the most used marker to visualize PNNs in histological analyses and one of the most studied protein of the neural extracellular matrix. A typical stained morphology of PNNs (Dickens et al., 2022), with this protein occasionally surrounding few PV^+^ cells (Figure 2C) was observed as early as DIV50, suggesting the formation of some early PNNs, mostly in SAG cultures.

**Figure 2.**
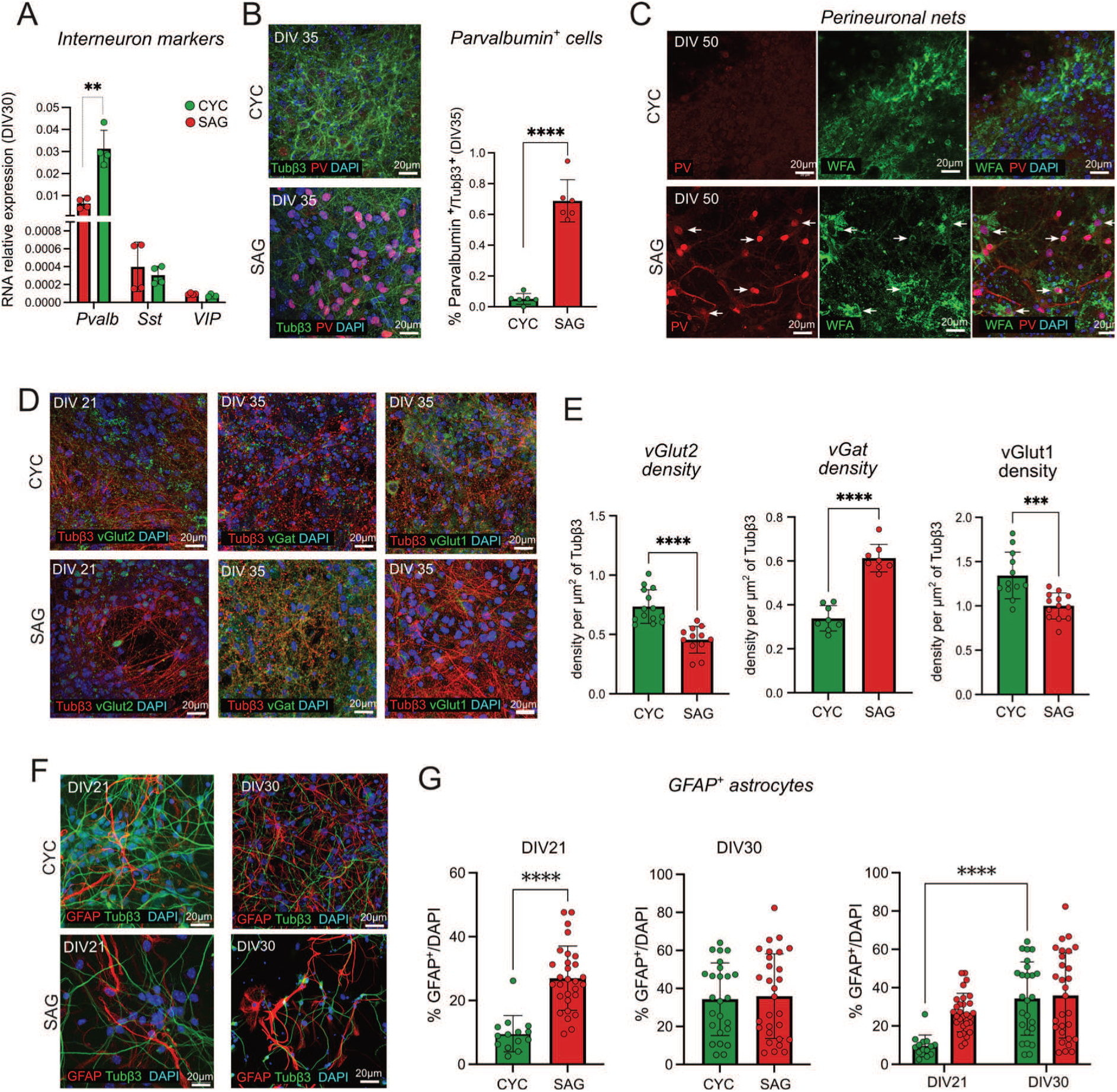
Expression of excitatory and inhibitory neural markers in CYC and SAG cultures. A) Relative expression of interneurons markers at DIV30 evaluated by qRT-PCR in the two different cultures (n = 4 independent experiments). Mean ± SD is shown; Multiple unpaired t-test with Holm-Šídák correction method. B) Representative confocal images of CYC and SAG cells stained with Tubβ3, Parvalbumin and DAPI at DIV35. Quantification of Parvalbumin (PV^+^) interneurons in CYC and SAG cells at DIV35. Mean ± SD is shown; Unpaired t-test; (n = 3 independent experiments). C) Representative confocal images of CYC and SAG cells stained with Perineuronal Nets markers: Parvalbumin, WFA and DAPI at DIV50. White arrows indicate the PNNs-like structures surrounding PV^+^ neurons. D) Representative confocal images of glutamatergic and GABAergic markers: vGlut2 (DIV21), vGat and vGlut1 (DIV35). E) Quantification of glutamatergic and GABAergic markers shown in (D). Synaptic vesicles were measured in terms of density of Tubβ3 positive area covered by vesicles. Comparisons showed statistical differences between the two treatments (n = 3 independent experiments; unpaired t-test). F) Representative confocal images with CYC and SAG neurons stained with GFAP and Tubβ3 antibodies at DIV21 and DIV30. G) Comparison of the percentage of GFAP positive cells at DIV21, DIV30 and over time between the two treatments; Mean ± SD is shown. Unpaired t-test was performed to compare samples at each time point, and two-way ANOVA followed by Šídák’s multiple comparison test was performed for comparison over time. P-values: *p-value < 0.05, **p-value < 0.01, ***p-value < 0.001, ****p-value < 0.0001.

The immunolabeling of distinct markers of excitatory (vGlut1, vGlut2) and inhibitory (Pvalb, vGat) cortical neurons showed a different contribution of the two types of cells in SAG and CYC cultures, confirming the prevalence of excitatory and inhibitory neurons in CYC and SAG cultures, respectively (Figures 2D). We investigated the early glutamatergic marker vGlut2 at DIV25: we evaluated the density of vesicles by observing the colocalization of vGlut2 puncta with Tubβ3*^+^* fibers, and found a significantly higher density of vGlut2^+^ vesicles in CYC neurons as compared to the SAG neurons (Figure 2E). Moreover, by counting the number of DAPI positive nuclei surrounded by vesicle transporters, we evaluated 70% of vGlut2^+^ cells in CYC cultures (Cao et al., 2017), while only 15% of them were present in SAG cultures (Kempf et al., 2021) (Figure S2A,B). We also analyzed the later glutamatergic marker vGlut1 and found a significantly higher density of vGlut1 positive puncta in CYC neurons as compared to SAG neurons (Figure 2E), with a 80% of CYC neurons surrounded by VGlut1^+^ vesicles (Figure S2C,D). Focusing on GABAergic markers, we analyzed the vGat density at DIV35 in our cultures, highlighting a significantly higher number of vGat^+^ vesicles in SAG neurons as compared to CYC neurons (Figure 2E).

To assess the percentage of astrocytes in CYC and SAG cultures we evaluated the presence of positive GFAP labeled cells at two time points of maturation, DIV21 and DIV30. We found a higher percentage of GFAP^+^ cells at DIV21 in SAG cultures as compared to CYC cultures, suggesting that SAG cultures complete neurogenesis and start astrogliogenesis earlier than CYC cultures. However, at DIV30 GFAP^+^ cells reached the same percentage (40%) in both cultures (Figure 2F,G), indicating that both have completed cell differentiation at this time.

Altogether, our observations indicate that timely treatment with either CYC or SAG molecules from DIV5 to DIV11 generated pallial NPCs with the competence to differentiate into neurons exhibiting a gene expression profile typical of dorsal and ventral telencephalic neurons, respectively. Moreover, CYC cells produced mostly glutamatergic pyramidal neurons while SAG cultures were enriched in GABAergic cells, mainly PV^+^ interneurons, similar to the inhibitory interneurons migrating to the embryonic developing cortex. Notably, both CYC and SAG cultures produced an optimal ratio of astrocytes (40%), which is anticipated to support functional network activity.

### Morphological analysis of CYC and SAG neurons in mixed cultures

The correct integration of ventral telencephalic neurons into the cortex is a regulated process requiring saltatory migration (Marín et al., 2010). We evaluated whether early SAG neurons can differentiate when mixed to isochronic CYC neurons and *vice versa*, in an *in vitro* environment and in the absence of regulated migration.We thus implemented mixed cultures with 80:20 and 50:50 (CYC:SAG) cells to model different ratios of excitatory and inhibitory neurons. We then analyzed the morphology of developing SAG or CYC labeled single neurons in isochronic cultures of unlabeled pure and mixed cells. To identify the soma and neurites of individual neurons, we transduced CYC or SAG cells with EGFP under the control of EF1ɑ promoter at DIV7 (Figure 3A) and mixed them at DIV17 in a 1:100 ratio with four types of unlabeled cultures: pure CYC and SAG cells, and 80:20 and 50:50 cultures (Figure 3B,C).

**Figure 3.**
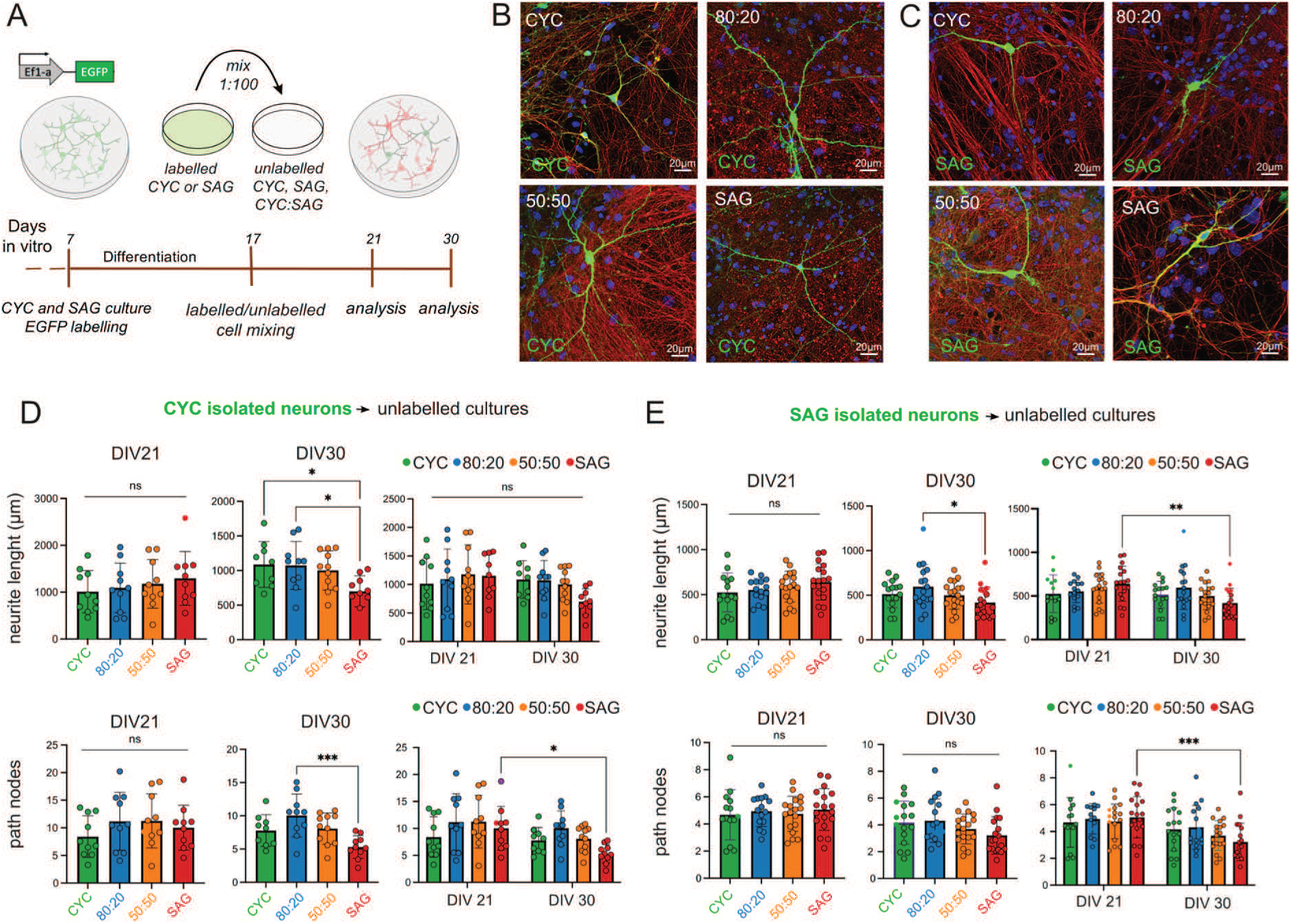
Morphological analysis of CYC and SAG neurons in mixed conditions. A) Experimental protocol: CYC and SAG progenitors were separately labeled at DIV7, differentiated to DIV17, then mixed and plated on different culture substrates. Immunofluorescence analysis was performed at DIV21 and DIV30. B-C) Representative confocal images of EGFP^+^-CYC neurons and EGFP^+^-SAG neurons, respectively, in the four different conditions: CYC, SAG, 80.20 and 50.50 mixed cultures. D-E) Quantitative analysis of neurite length and branching patterns (number of nodes) of CYC and SAG neurons, respectively, in all four conditions. Comparisons were done at DIV21, DIV30 and then, the two time points were also compared; ; Mean ± SD is shown. Ordinary one-way ANOVA with Tukey’s multiple comparisons test was performed to compare samples at each time point, and two-way ANOVA followed by Šídák’s multiple comparisons test was performed for comparisons over time. P-values: *p-value < 0.05, **p-value < 0.01, ***p-value < 0.001, ****p-value < 0.0001, ns = not significant.

We performed the analysis at two time points, DIV21 and at DIV30, to capture early neural development and the onset of neural maturation. We examined specific parameters to evaluate the morphological complexity of these neurons, such as the overall length of neurites, branching patterns (number of nodes within neurites), and the percentage of dendritic spines. We observed a decrease in neurite length of CYC neurons in SAG cultures at DIV30, although this reduction is not maintained over time when comparing the two time points. We also found a reduced number of their nodes in the SAG culture environment as compared with the other ones, especially with 80:20 condition, resulting in a lower maturation of these neurons over time as well (Figure 3D). Nonetheless, we found a similar significant decrease in the length of neurites of SAG neurons in the SAG cultures at DIV30, compared with those in 80:20 cultures, and over time. In addition, a reduced number of nodes was observed only when comparing the two time points, concluding that SAG neurons also showed an altered maturation in branching patterns (Figure 3E).

The analysis of the percentage of the dendritic spines (Figure S3A,B) highlighted a reduced percentage of spines of CYC neurons in 50.50 and SAG conditions, both at DIV21 and DIV30 (Figure S3C). Accordingly, comparison of the two time points suggests that CYC neurons generate more spines under homotypic conditions, with an increased percentage of spines in the CYC and 80:20 configurations (Figure S3C). On the other hand, the maturation of SAG neurons seem to be significantly impaired at DIV21 in the mostly inhibitory environments, while there are no differences at DIV30 between the four conditions (Figure S3D). Despite this, comparing both time points, they show an increasing percentage of spines only in 50:50 and SAG conditions (Figure S3D), indicating a maturation step that is in line with the knowledge that medium spiny neurons in the striatum form abundant dendritic spines during development (Steiner, H. and Tseng, K.-Y., 2017). Overall, our results indicate that the composition of the surrounding neuronal population might influence the morphological maturation of a neuron. Specifically, although neuron maturation is not impaired in heterotypic environments, we observed alterations in neurite length, branching complexity, and dendritic spine density in both CYC and SAG neurons when cultured in mostly inhibitory environments. Thus, these morphological variations suggest that the diverse cellular environments may impact the functional integration and connectivity of these neurons within cortical circuits.

### Functional activity development of pure and mixed CYC and SAG neuronal networks

Primary cortical neurons cultured in adhesion self-organize and develop neuronal networks displaying various activity patterns (Charlesworth et al., 2015; Dias et al., 2021). In particular, a balance between excitatory and inhibitory neurons forms the basis for functional neural networks, which is crucial for normal cognition and memory. This balance is maintained at the level of individual neurons by an appropriate ratio of excitatory to inhibitory synaptic inputs, and at the network level by regulating the interaction between various excitatory and inhibitory circuits (He and Cline, 2019). Therefore, we decided to analyze pure CYC cultures (simulating cortical development without the contribution of ventrally migrated interneurons), SAG cultures (modelling the striatal network) and mixed CYC:SAG cultures with different ratios of excitatory neurons and PV^+^ inhibitory interneurons, in order to simulate various E/I conditions.

Using a high-density microelectrode array (HD-MEA) comprising 4096 channels, we conducted a longitudinal study of various activity parameters between pure CYC and SAG cultures and mixed cultures with an 80:20 or 50:50 ratio. As CYC and SAG cultures contained 7% and 73% of PV^+^ cells (Figure 2B), the theoretical PV^+^ cells in 80:20 and 50:50 cultures is 20.2% and 40% in 80:20 and 50:50, respectively (Figure 4A,C, Supplementary Movies S1-8). The pure CYC network developed differently than SAG cultures: cells tended to cluster (Figure S4A), although the average cell density was comparable between the two types of culture (Figure S4B). Moreover, the thickness of CYC cultures was significantly larger than that of SAG cultures (Figure S4C), suggesting different adhesion properties and maturation timing of connectivity between the two culture types. This uneven distribution of CYC cells on the MEA surface was expected to decrease the ratio of efficient recording electrodes. Accordingly, the number of active channels observed at DIV45 in CYC cultures (n=500 ± 100) was lower compared to 80:20, 50:50 and SAG cultures (n=1600 ± 400, n=1700 ± 300, n=1174 ± 35, respectively) (Figure S4D).

**Figure 4.**
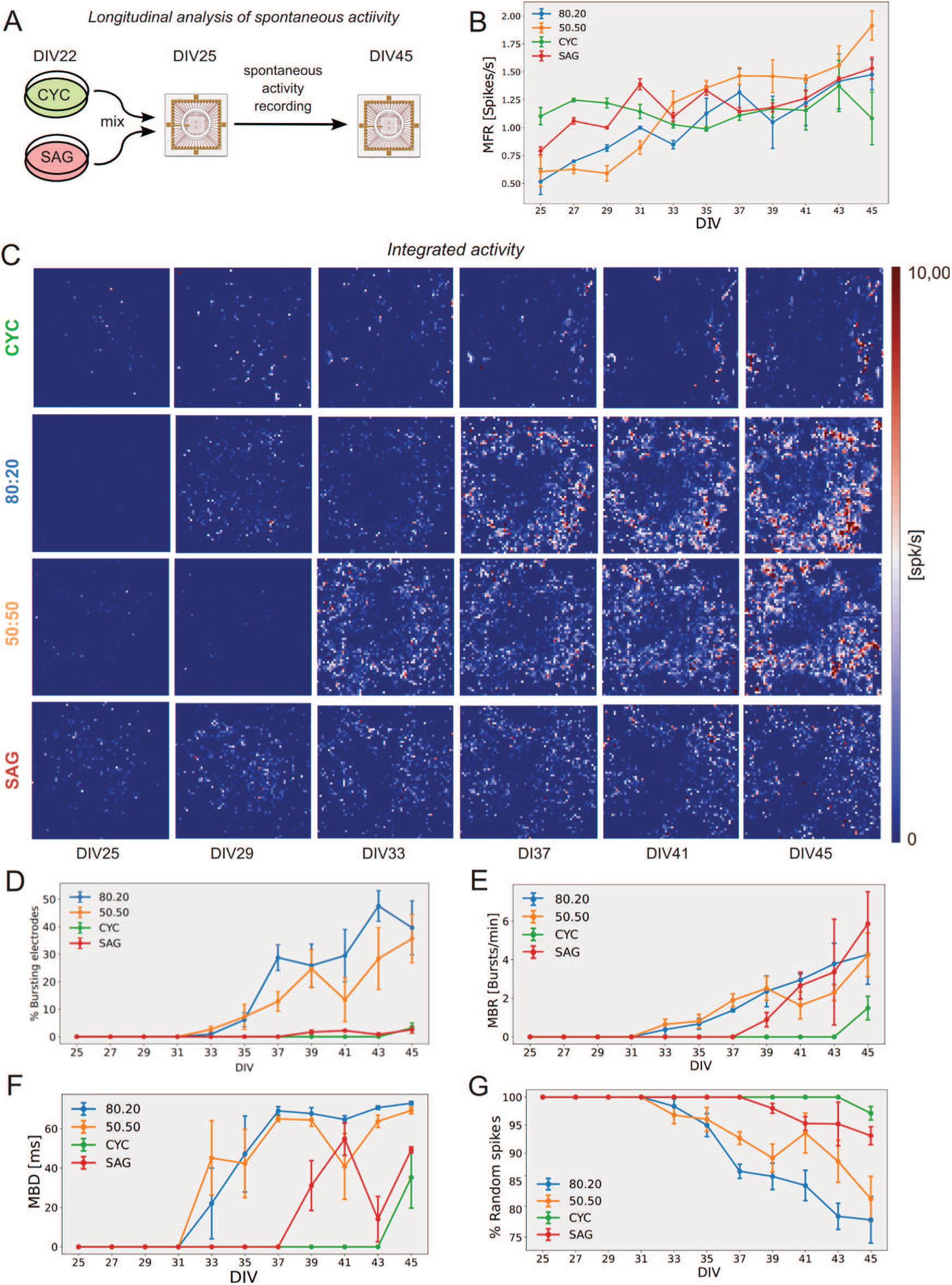
Longitudinal analysis of electrophysiological activity of CYC, SAG and mixed cultures. A) Outline of the longitudinal study of neural activity. B) Mean firing rate (MFR) of individual channels over time (n = 3 independent experiments; Mean ± SEM is shown). C) Representative global HD-MEA heatmap activity of the four conditions in different time points, depicting the firing rate (spk/s) of the electrophysiological activity; each pixel represents one channel. D, E, F, G) Percentage of bursting electrodes, Mean Burst Rate (MBR), Mean Burst Duration (MBD) and percentage of random spikes of individual channels (n = 3 independent experiments; Mean ± SEM is shown).

For each culture, we first analyzed the mean firing rate per channel (MFR), the mean burst rate (MBR) and duration (MBD), the percentage of bursting electrodes and of random spikes. From DIV25 to DIV45 the MFR of all the cultures increased over the time of analysis with no significant differences between cultures (Figure 4B, C). A clear difference between pure and mixed populations was observed in the number of bursting electrodes, which was much lower in pure populations (Figure 4D, Supplementary movies VS5-8). Spike bursts (see Methods for definition) appeared first in 50:50 and 80:20 cultures (DIV33), preceding SAG cultures (DIV39) and CYC cultures (DIV45) (Figures 4E,F). MBR reached higher values in 80:20, 50:50 and SAG cultures compared to CYC cultures (Figure 4E). MBD was higher in 80:20 and 50:50 cultures compared to SAG and CYC cultures (Figure 4F). However, although the MBR in CYC cultures was much lower than that in the SAG condition, the MBD of both was comparable, suggesting that a minimum number of spikes is intrinsic to the burst generation mechanism. Overall, these observations indicate that pure CYC and SAG populations are less inclined to generate burst activity as compared to mixed populations. This result, together with the observation that pure populations yield the highest percentage of random spikes (Figure 4G), while showing a comparable MFR to mixed populations (Figure 4B), indicates that mixing of the two cell types greatly enhances the capability to generate bursting activity.

### Pure and mixed CYC and SAG cultures show distinct patterns of network burst activity

Spontaneous network activity underpins the development of functional networks in early stages (Teppola et al., 2019). A hallmark of this activity is the recurrent occurrence of intense, time-constrained network bursts (NBs) that rapidly propagate throughout the entire dissociated culture *in vitro* (Okujeni et al., 2017; Weihberger et al., 2013). We analyzed NBs in our cultures as defined by the peaks in the firing rate (see Methods for further details). NBs were generated first in CYC:SAG mixed cultures starting from DIV31 and then in pure SAG cultures, although at a very low extent, from DIV35, whereas they were almost absent in pure CYC cultures (Figure 5A,B). In mature cultures at DIV45, NB duration (NBD) was comparable in the four cultures (Figure 5C), indicating that this parameter is independent of the NBs frequency. Although CYC and 80.20 cultures have the common presence of a low percentage of PV^+^ interneurons and high percentage of vGlut1^+^ neurons, these cultures showed an opposite trend in generating NB. Therefore, we hypothesized that they developed a different degree of excitatory/inhibitory connectivity.

**Figure 5.**
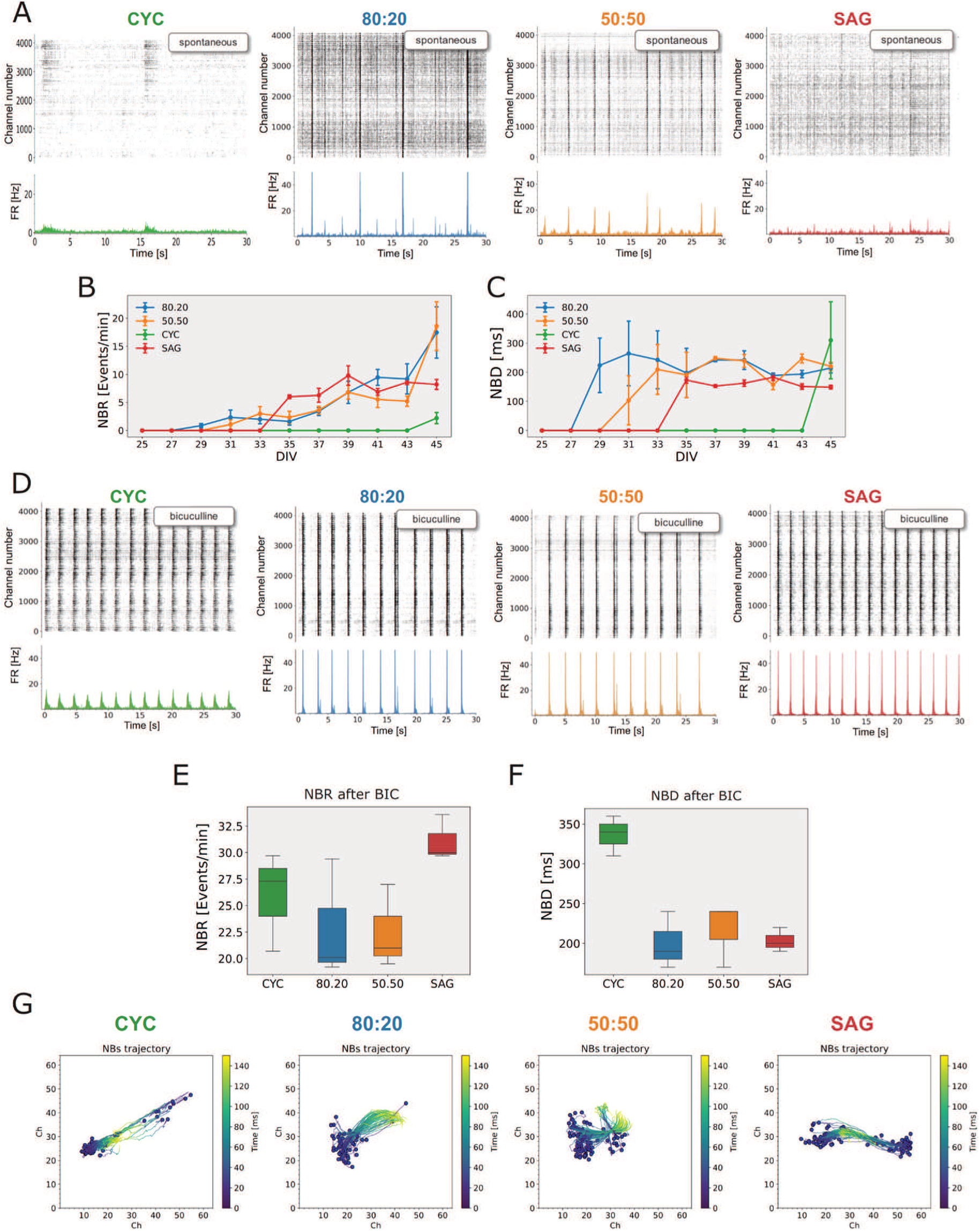
GABA inhibition critically affects network activity in pure and mixed CYC and SAG cultures. A) On the top: representative raster plot of activity (spike times per channel) for each culture condition at the analysis endpoint (DIV45); on the bottom: average network firing rate (number of spikes per channel per time window) computed in 20 ms time bin. B,C) Network Burst Rate (NBR) and Network Burst Duration (NBD) (n = 3 independent experiments; Mean ± SEM is shown). D) Raster plots of network activity after Bicuculline administration. E,F) Boxplots reporting the Network Burst Rate (E) and Network Burst Duration (F) after Bicuculline administration in all four conditions (n = 3 replicates). G) Analysis of the center of activity trajectories (CATs) in the four conditions. Each blue dot represents the physical center of mass of the activity where the NB starts, while the colored line represents its own trajectory. The colored scale bar represents the time elapsed during the propagation of a network burst (NB).

We thus investigated the role of GABA, AMPA and NMDA receptors in NB occurrence of mature networks (DIV45). AMPA and NMDA receptors play a key role in maintaining NB activity (Jimbo et al., 2000) and, consistently, we found that their antagonists, CNQX and AP5, respectively inhibited and blocked NBs in all the cultures (Figure S5A-D), changing significantly their firing activity (Figure S5E).

As the four cultures have different degrees of GABAergic signaling, mainly due to the presence of different percentages of PV^+^ interneurons (compatible with theoretical expected values, see Figure S5F,G), they respond differently to the administration of Bicuculline (BIC), a GABA_A_ receptor antagonist. In particular, compared with the spontaneous NBR at DIV45, 80:20 and 50:50 (CYC:SAG) cultures slightly increased the rate of their NBs after BIC administration (Figure 5D-E). Pure CYC and SAG cultures showed a massive increase of NB rate and duration. Notably, CYC NBR was even higher than the 80:20 one. This effect was unexpected, as 80:20 cultures contain more PV^+^ cells than CYC cultures (Figure S5G). Moreover, the NBD of the CYC culture was much longer than that of the other three cultures after BIC administration (Figure 5F). These observations indicate that the PV^+^ cell ratio does not correlate with the NB activity. In fact, very low PV^+^ cell ratio strongly inhibits NBR compared to cultures with intermediate levels of PV^+^ cell ratio. We anticipate that this might be due to the development of different patterns of structural and functional connectivity of the four types of cultures, although this hypothesis requires further experimental investigation.

To quantify the propagation of NBs after bicuculline administration, we used Center of Activity Trajectory (CAT) analysis (Chao et al., 2007). This method determines the spatial and the time evolution of the electrical activity of each neural culture during network burst events, determining a sort of center of mass for the spikes, where for each channel the mass is replaced by the number of recorded spikes. CAT analysis (Figure 5H) showed that NBs propagated with similar properties in the four cultures when the inhibitory activity was suppressed.

In conclusion, BIC appeared to unmask an intrinsic capacity of CYC and SAG cultures to generate and propagate NBs.

### GABAergic and glutamatergic components differently impact on pure and mixed CYC and SAG functional networks

As GABAergic inhibition is responsible for masking the intrinsic NB activity of pure cultures compared to mixed cultures, we investigated whether a specific functional connectivity analysis could explain the different effect of BIC on the NB activity of the four conditions. We thus compared the functional network connectivity, calculated as the correlation of activity between channels (see methods for a detailed description), before and after the administration of BIC (Figure 6A) and, for comparison, of CNQX and AP5 (Figure S5). The correlation analysis allowed us to calculate the number of nodes and links of the spontaneous network activity (Figure 6B) and their changes after drug administration (Figure 6C). In the absence of drugs, the mixed cultures showed a similar number of nodes and links while SAG and CYC networks had fewer, with CYC showing the lowest number (Figure 6B). The administration of each of the three drugs differently affected the four cultures (Figure 6C, Figure S6). The most striking effect was the large increase of nodes and links exerted by BIC in CYC condition, exceeding the increase observed in SAG condition. This is noteworthy considering that SAG cultures had a much higher proportion of PV^+^ neurons and lower proportion of glutamatergic vGlut1^+^ neurons than CYC cultures and suggests that a very complex part (nodes and links) of the CYC functional network was under GABAergic inhibition. Indeed, some studies have reported that the excitatory activity of pyramidal neurons in the cerebral cortex can be significantly influenced by PV^+^ interneurons in order to compensate for the overactive excitatory force (Haider et al., 2006; Xue et al., 2014). Finally, while CNQX had almost no effect on SAG and 50:50 cultures as compared to CYC and 80:20, AP5 decreased the number of nodes and links in all the cultures, although at different extents (Figure 6C). This indicates that the activity of NMDA receptors is important in the developmental regulation of synaptic transmission, also mediated by both AMPA and GABA_A_ receptors (Lu et al., 2011; Marsden et al., 2007), and plays a pivotal role in establishing NBs activity and maintaining the delicate E/I balance.

**Figure 6.**
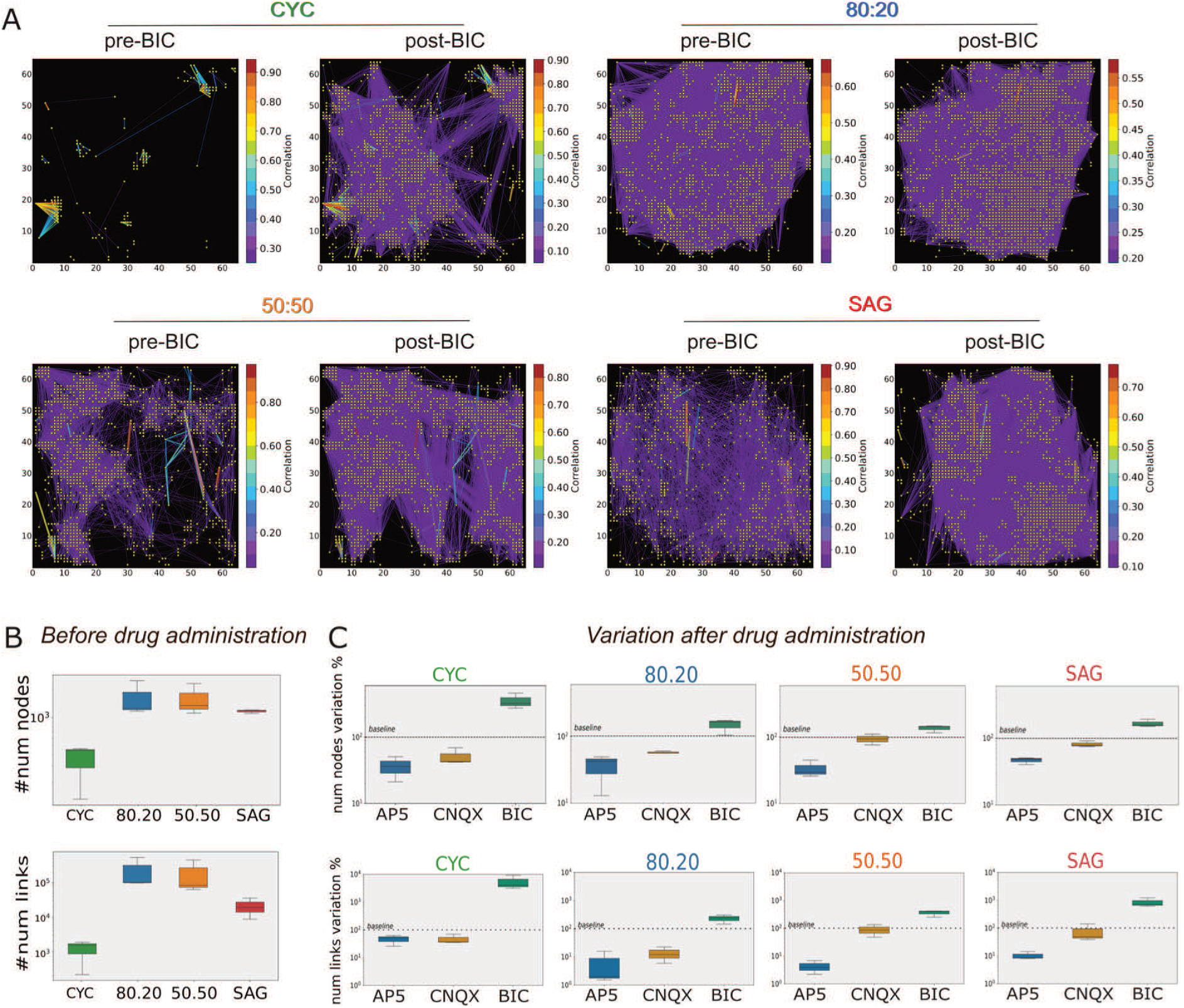
GABA inhibition unmasks complex connectivity of the CYC network. A) Connectivity plots of representative neuronal cultures during spontaneous activity before (pre-BIC) and after (post-BIC) bicuculline administration. Each yellow point represents a node of the functional graph; colored lines represent the correlation strength between two points (only the 10% of the functional links are shown). The color bar indicates the correlation index. B) Number of nodes and links of the obtained connectivity graphs for all cultures before drug administration (n = 3 replicates). C) Number of nodes and links of the obtained connectivity graphs for all cultures after drugs administration (n = 3 replicates).

### Network stimulation discloses different signal propagation properties of pure and mixed cultures

MEA technology has been proved to be a very important tool to explore the impact of electrical stimulation in different types of cell cultures (Parodi et al., 2023; Scarsi et al., 2017). Thus, we analyzed the patterns of activity evoked by electrical stimulation in our cultures and compared them to the patterns of spontaneous activity. At DIV50, we stimulated a single channel with 25 biphasic pulses of 10µA current with a duration of 100 µs, at a frequency of 0.1 Hz. For each electrode, the response to the stimulus was measured by comparing spike counts before and after the pulse in a specific time window. To account for variability, the stimulus was repeated 25 times, responses were averaged and confidence intervals for the average response were calculated (see Methods for further details).

The single channel stimulation evoked the activity of channels located at different distances within a 5ms delay, suggesting direct functional connectivity between the stimulated channel and the responding ones (Figure 7A, Figure S7A-D). The temporal pattern of stimulus propagation was different in the four cultures. The 80:20 condition was the most responsive to the single channel stimulation (Figure 7A). In this condition, the stimulus first recruited a high number of positively correlated channels by increasing their spiking activity within 30 ms (red channels in Figure S7A), then evoked activity silencing in negatively correlated channels with a delay of 60 ms (yellow channels in Figure S7A). In 50:50 mix and SAG cultures the induction of positively correlated active channels was moderate, while the silencing of negatively correlated channels was more pronounced and earlier compared to 80:20 cultures (Figure S7B,D). CYC cultures behaved differently, showing recruitment of almost all activated channels within 5 ms and silencing of negatively correlated channels with a delay of 60 ms, more like the 80:20 mix (Figure S7C). The longitudinal analysis of the activity-related channels also allowed us to evaluate the spatial dispersion of the signal, as a function of their physical distance from the stimulation point (Figure 7B, see Methods - Dispersion index - for detailed explanation).

**Figure 7.**
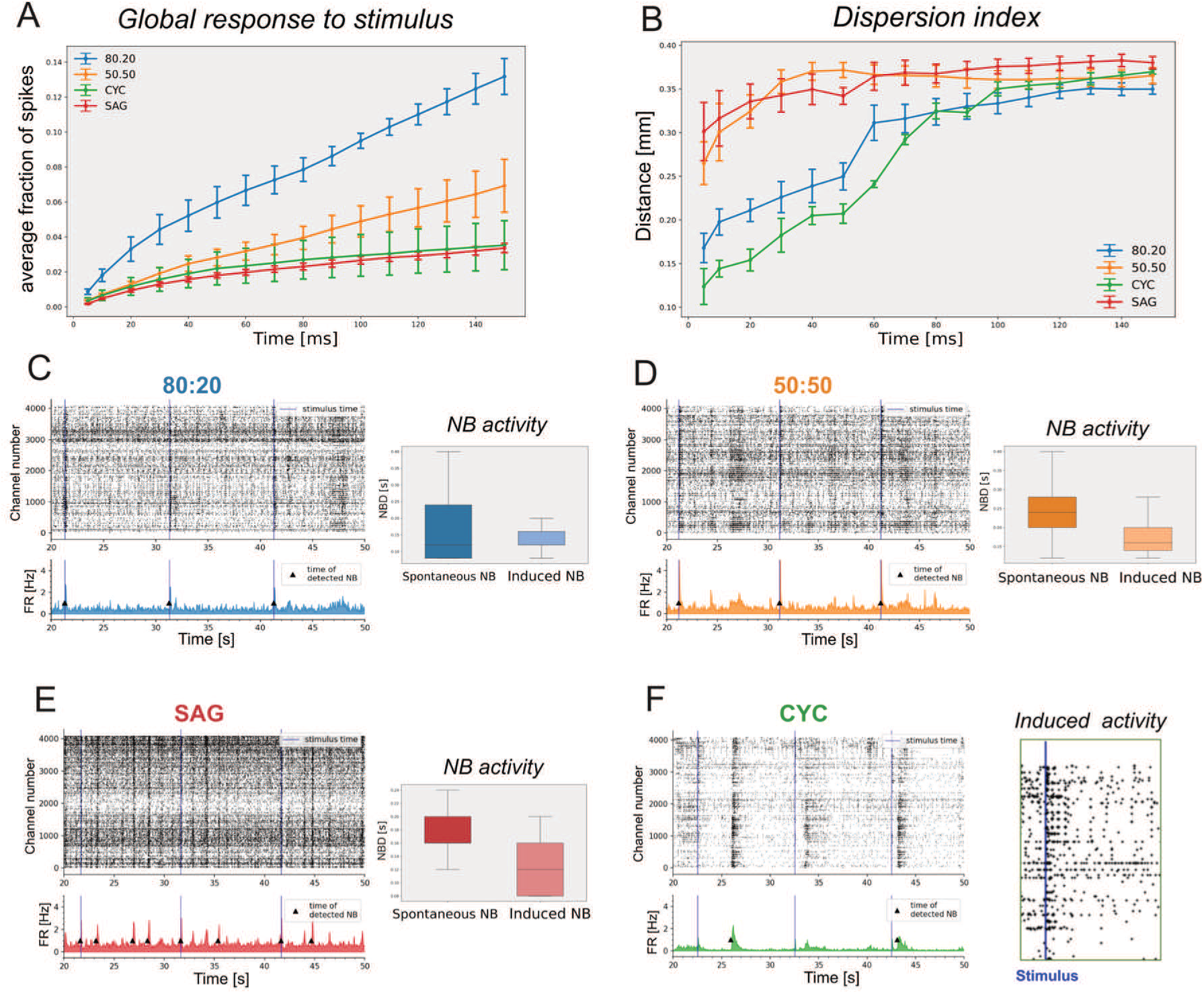
Analysis of global response, dispersion index and network bursts induction evoked by stimulus are affected by E/I ratio. A) Distribution of the global response of the four conditions to the single channel stimulation, showing the average fraction of spikes starting from 5 ms to 150 ms (n = 3 independent experiments; Mean ± SEM is shown). B) Dispersion index indicating the distance of responses from the single channel stimulation point in 150 ms (n = 3 biological replicates; Mean ± SEM is shown). C-F) Representative raster plots showing the stimulus time (blue line) and the detected NBs in all four conditions. On the right in F: enlarged detail of CYC raster plot, highlighting only the electrodes that show a response to the stimulus. Boxplots represent the Network Burst Duration (NBD) of the spontaneous NBs and of the induced ones (n = 3 replicates).

Our findings reveal a correlation between the dispersion index of the signal evoked by the stimulus and the PV^+^ cell ratio in the early phase following stimulation, with all four cultures exhibiting a common peak in dispersion index of approximately 0.3-0.4 mm at 150 ms post-stimulation. 80:20 cultures show the most robust global stimulus response but low dispersion index at early times, with activity observed in channels very close to the stimulation site. This observation may indicate a tendency for these cultures to form local circuits. Conversely, while exhibiting more distant initial responses from the stimulated channels, 50:50 and SAG cultures demonstrate a significantly lower overall global response. Therefore, we hypothesize that stronger inhibition in these cultures may locally suppress the response to the initial stimulus by promoting it farther from the stimulation site through long-range neural projection processes. We also concluded that intermediate PV^+^ cell ratios support the best global activity due to a compromise between signal spatial propagation and signal inhibition.

Notably, single channel stimulation was never able to propagate to the entire network and to generate NBs. Thus, we speculated that the stimulation of a proper number of spontaneously active electrodes in cultures could evoke NBs. We selected the channels with the highest firing rate (see methods for further details) for simultaneous stimulation, optimizing a number of channels in each culture (n = 7) and applying single stimuli with the same parameters of the single channel stimulation.

In 80:20, 50:50 and SAG conditions, the electrical stimulation evoked a synchronized network activity aligned with the emission of the stimulus and comparable to spontaneous NBs (Figure 7C-E). CYC cultures did not generate NBs in response to the stimulus (Figure 7F): indeed, we found that the response to stimuli did not propagate, activating only specific channels of the plate (enlarged detail of the raster plot in Figure 7F). This last observation suggests that a proper inhibitory ratio is required to integrate and organize a network circuit capable of propagating NBs (Tremblay et al., 2016). Moreover, in all the other three configurations, we evaluated the duration of the induced network burst (NBD). In particular, in the 80:20 configuration (Figure 7C), we observed that the duration of the electrical induced NBs was similar to the one of spontaneous NBs, leaving the network almost unchanged. On the other hand, in both 50:50 and SAG conditions, the duration of the induced NBs was shorter as compared to the spontaneous ones (Figure 7D,E). Moreover, the electrical induced NBs had also a reduced duration as compared to the ones induced by chemical stimulation with Bicuculline (Figure 5G), which, on the contrary, are more similar to spontaneous NBs. These findings demonstrate that the presence of PV^+^ interneurons significantly impacts the dispersion of neuronal activity, influencing network connectivity and determining the capacity of cortical networks to generate synchronous network bursts.

## DISCUSSION

We modeled the development and maturation of cortical and striatal cells *in vitro,* focusing on their capability to initiate and sustain functional network activity in isolated and mixed cultures. Indeed, we found that neuronal cultures with different molecular identities developed distinct network activity.

Analysis of our CYC and SAG cultures indicated that they developed expression of key marker genes consistent with cortical and striatal identity and matured opposite ratios of PV^+^ inhibitory interneurons, very high (73%) in SAG cultures and very low (7%) in CYC cultures. Moreover, CYC cultures were enriched in glutamatergic vGlut1^+^ neurons (73%) whereas vGlut1^+^ neurons were 16% in SAG cultures. We observed that isolated CYC and SAG neurons could mature in each other’s cultures, allowing us to implement mixed cultures carrying different proportions of PV^+^ GABAergic interneurons. This was fundamental to investigating the activity properties that emerge in networks with different amounts of inhibitory activity. Our approach aimed to understand some of the basic requirements for the generation of neuronal networks capable of spontaneous correlated activity similar to that recorded in early encephalic regions (Chiu and Weliky, 2001; Corlew et al., 2004; Crochet et al., 2005; Harsch and Robinson, 2000). The main constraint of *in vitro* 2D modeling of cortical networks is the difficulty to reproduce the migration of ventral telencephalic interneurons in a culture dish. However, the electrophysiological recording of CYC cultures allowed us to model how cortical network activity develops without migrated interneurons. Moreover, comparing CYC neuronal culture activity with that of CYC and SAG cells mixed up in 80:20 and 50:50 ratios, or pure SAG cultures, highlighted unexpected effects of PV^+^ GABAergic neurons on spontaneous activity patterns.

We found that, although the four types of cultures develop a similar MFR, pure CYC cultures showed a very low correlated (burst) activity. Nonetheless, upon the addition of a low percentage of SAG neurons (20%), the activity of 80:20 cultures changed dramatically, exhibiting more synchronous firing patterns and reducing the percentage of recorded random spikes, while raising that of bursting electrodes. Moreover, the 80:20 cultures showed a significant increase of their ability to generate spontaneous correlated activity. In fact, their activity improved significantly in terms of network burst rate (NBR) compared to the very low activity observed in pure CYC cultures, reaching levels comparable to other types of cultures. In addition, the 80:20 cultures also showed the highest network burst duration (NBD). This is a puzzling observation, considering that SAG cultures contain 73% of PV^+^ GABAergic neurons and, when cultured alone, showed almost no network bursting activity. Thus, we speculated that a proper ratio of GABAergic neurons is required to mature local circuitry capable of burst activity formation and spreading. Concerning this aspect, we may hypothesize that low PV^+^ cell ratio might favour a higher number of inhibitory synapses to glutamatergic neurons, which would reduce the NB activity, while intermediate PV^+^ cell ratios may increase the probability of the formation of synapses between PV^+^ neurons, supporting network activity.

The presence of different circuits in the four culture types is suggested by the different types of network activity generated under spontaneous conditions, upon GABAergic release by BIC administration or electrical stimulation. CYC cultures, which showed the lowest spontaneous NBR, generated the highest NBR together with SAG cultures after bicuculline treatment, but were unable to induce NBs upon electrical stimulation. These observations, together with the shortest dispersion index shown by CYC cultures, point out that, unexpectedly, a too high E/I ratio develops a different network circuitry, confirming that an optimal E/I ratio is required to form local inhibitory networks capable of developing and spreading correlated activity.

## EXPERIMENTAL PROCEDURES

Mouse embryonic stem cells (ESCs) were differentiated into cortical (dorsal telencephalic) or striatal (ventral telencephalic) lineages following a four-step protocol. In the first step (DIV0-DIV5) cells were cultured in a chemically defined minimal medium supplemented with Wnt and BMP inhibitors (WiBi) and plated on poly-ornithine/laminin-coated (PL) surfaces. In the second step (DIV6-DIV10) cortical and striatal cells were generated adding Cyclopamine (Sigma, S-4116, 3μM) and SAG (Santa Cruz Biotechnology, SC-212905, 0.1μM), respectively, to the WiBi medium. In the third step (DIV11-DIV20), cells were replated onto PL and maintained in Neurobasal-A. From DIV20 onward, neurons were maintained in Neurobasal-A medium supplemented with 0.2mM Ascorbic Acid and 20 ng/ml recombinant human BDNF protein. To prepare the mixed cultures of 80:20 and 50:50, neurons were splitted at specific time points. For MEA preparation, Cyclopamine-treated and SAG-treated neurons were detached at DIV22, counted, mixed with the designed proportions and seeded on sterilized high-density microelectrode array (HD-MEA Accura, 3Brain) (60000 cells/chip) and allowed to adhere O/N.

Quantification analyses for density of glutamatergic and GABAergic vesicles were done by using the ImageJ Synapse Counter plugin (https://github.com/SynPuCo/SynapseCounter). To study neuronal development in pure and mixed cultures, Cyclopamine- and SAG-treated cells were transduced with EGFP lentivirus at DIV7 and selected via PuroR. At DIV17, EGFP-labeled and unlabeled neurons were combined to create pure (Cyclopamine-only, SAG-only) and mixed (80:20, 50:50) cultures, each with 1% EGFP-positive cells, and seeded on PL-treated glass (150,000 cells/cm²). Cultures were fixed with 2% PFA at DIV21 and DIV30 and analyzed by immunofluorescence. Neural length and the node number were evaluated by the FIJI SNT plugin.

Electrophysiological recordings were conducted using high-density CMOS-based 4096 microelectrode arrays (Accura, 3Brain) to evaluate spiking activity, bursting behavior, and network synchronization. Drug responses and functional connectivity were analyzed using custom Python scripts, with spike sorting and network analysis performed using principal component analysis (PCA) and cross-correlation methods. The response to stimuli was measured by comparing spike counts before and after the stimulus within a defined time window (5 to 150 ms). The stimulus was repeated 25 times, responses were averaged and statistical confidence intervals were calculated. For data visualization, responses were displayed on a grid, with pixel intensity indicating response strength. The dispersion index, quantifying how spread out significant responses were across the grid, was calculated estimating the average distances between responsive electrodes.

Comprehensive descriptions of all experimental procedures are provided in the Methods section of the supplemental information.

## Resource Availability

### Lead contact

Further information and requests for resources and reagents should be directed to and will be fulfilled by the lead contact, Federico Cremisi.

### Materials availability

This study did not generate new unique reagents. However, any questions about reagents or animals used can be directed to the lead contact.

## Acknowledgements

We are thankful to Prof. Robert Vignali and Dr. Lucio Calcagnile for helpful discussions and to Dr. Maria Antonietta Calvello, Dr. Vania Liverani and Mr. Alessandro Puntoni for technical support. We thank Mr. Stefano Guglielmo, Dr. Claudia Alia and Dr. Nicola Origlia for advice on the functional analysis of cultured networks. The research was supported by the Matteo Caleo Foundation, by intramural funding of IIT (S.G. and L.P.) and Scuola Normale Superiore (FC), by the PRIN grant #2022M95RC7 from the Italian Ministry of University and Research (FC) and by the Tuscany Health Ecosystem - THE grant from MUR (FC, AD).

## Author Contributions

E.C. and F.C. conceptualized and designed the study and wrote the manuscript. E.C. F.T. and L.I. performed the experiments. E.C. set up the strategy of culture treatments, the connectivity analysis and the longitudinal functional study. F.T. set up MEA stimulation. L.P. conceptualized the experiments of gene expression analysis. L.I. set up the time-series analysis of MEA data under the advice of A.D. G.L performed the analysis of MEA stimulation under the supervision of G.A. All authors discussed the results and commented on the manuscript.

## Methods

### Maintenance of mouse ES cells

Mouse embryonic stem cells E14Tg2A were expanded and differentiated as follows (Bertacchi *et al*., 2015). Cells were kept on 0,1% gelatin-coated culture dishes, seeding at a density of 50000 cells/cm^2^ and splitting when at 70-80% of confluence. Cells were maintained in ES cell medium based on GMEM (Gibco, 11710035), containing 10% Fetal Bovine Serum (Euroclone ECS0180L), 2mM Glutamine, 1 mM Sodium Pyruvate, 100 U/ml Penicillin-streptomycin, 1mM Non-essential amino acids, 0.05mM β-mercaptoethanol. The medium was replaced daily.

ESCs were expanded in ES medium for three to four passages, then the cells were cultured in 2i+LIF medium, based on GMEM supplemented with 1x N-2 Supplement (Gibco, 17502001), 1x B-27 Supplement minus Vitamin A (Gibco, 12587010), 2mM Glutamine, 1mM Sodium Pyruvate, 1mM NEAA, 0.05mM β-mercaptoethanol, 1μM MEK inhibitor PD0325901 (Mirdametinib, Selleck Chemicals, S1036), 3μM GSK3 inhibitor CHIR99021 (Sigma-Aldrich, SML1046), and 10ng/mL recombinant mouse LIF(Silva et al., 2008).

### Differentiation of ES cells to different cortical fates

Differentiation of ES cells into cortical neurons was performed as previously described(Bertacchi et al., 2015), (Tonelli et al., 2025). For neural induction, a chemically defined minimal medium (CDMM) containing DMEM/F12 (Gibco, 11320033), 2mM Glutamine, 1mM Sodium Pyruvate, 0.1mM NEAA, 0.05mM β-mercaptoethanol, 1x N-2 Supplement, and 1x B-27 Supplement minus Vitamin A was used. The differentiation protocol started by culturing mESCs (3x10^4^ cell/cm^2^) onto 0,1% gelatin plastic coated dishes in 2i+LIF Medium for one day, marked as Day *in Vitro* -1 (DIV -1).

The next day (DIV0), the medium was replaced with Wnt and BMP double inhibition (WiBi) medium: CDMM with 2.5μM 53AH (Wnt pathway inhibitor, Cellagen Technology, C5324-2s) and 0.25μM LDN193189 hydrochloride (Bmp inhibitor, Sigma-Aldrich, SML0559). Cells were cultured in WiBi medium for 3 days (DIV0 - DIV3). On DIV3, differentiating ES cells were dissociated and seeded (30000 cells/cm2) in CDMM on dishes coated with Poly-ornithine (PLO, Sigma; 4 μg/cm2 in sterile water, 3 hours coating at 37°C) and purified mouse Laminin (msLam, Sigma-Aldrich, CC095-M; 1 μg/cm2 in PBS, O/N coating at 37°C). Next day, the medium was changed to WiBi medium and cells were cultured until DIV5 with daily medium changes.

From DIV5 until DIV10 chemicals as Cyclopamine (Sigma, S-4116, 3μM) and Smoothened agonist SAG (Santa Cruz Biotechnology, SC-212905, 0.1μM) was added to the WiBi medium to differentiate the two neural populations. At DIV7 cells were split again into mouse-Laminin coating dishes with a density of 110.000 cells/cm^2^.

At DIV11 until DIV21 cells were maintained in “young” Neurobasal (yNb), containing Neurobasal medium, 2mM Glutamine, 1mM sodium Pyruvate, 0.05mM β-mercaptoethanol, 0.2mM Ascorbic Acid (Vit. C), and B-27 Supplement minus Vitamin A 50x (NEAA were removed from the CDMM to prevent glutamate-induced excitotoxicity). From DIV13, half of the eNb medium was changed daily to allow conditioning of the medium by the differentiating neurons.

At DIV21, medium was changed to “old” Neurobasal medium (oNb), containing Neurobasal-A (Gibco, 10888022), 2mM Glutamine, 1mM sodium Pyruvate, 0.05mM β-mercaptoethanol, 0.2mM Ascorbic Acid (Vit. C), B-27 Supplement 50x (Gibco, 17504044), and 20 ng/ml recombinant human BDNF protein (Novus Biologicals, NBP2-52006). Starting the next day, oNb medium was changed every 2 to 3 days to condition the medium and allow the differentiated neurons to mature.

### Passaging and long-term culture of neurons

Usually, cells were passaged during differentiation to avoid overgrowth and hypoxia within large cell clusters, to improve immunofluorescence confocal imaging, and for electrophysiological recordings. For this purpose, cells were first washed with 1x Versene and then incubated with 1x Trypsin for 5-20 min at 37°C. When most of the cells detached, trypsin was inactivated by adding 20% FCS to the cell suspension and diluting in warm PBS (1:5 ratio).

During the last period of neuralization, the cell pellet was rinsed with a warm yNb medium containing 4μM of Rock inhibitor Y-27632 (Cell Guidance Systems, SM02) to reduce the mortality of post-mitotic neurons. Cells were seeded on PLO/msLam-treated glass (200000 cells/cm2) or sterilized high-density microelectrode array (HD-MEA Accura, 3Brain) (60000 cells/chip) and allowed to adhere O/N. The next day, the medium was changed, and the Rock inhibitor was removed. Occasionally, 1.5μg/mL msLam was added to the medium to increase neuron attachment and long-range projection.

### Preparation of 80.20 and 50.50 mixed cultured

To prepare the mixed cultures of 80.20 and 50.50, neurons were splitted at specific time points. For 3Brain Chip preparation, Cyclopamine-treated and SAG-treated neurons were detached at DIV22, counted, mixed with the designed proportions and seeded on sterilized high-density microelectrode array (HD-MEA Accura, 3Brain) (60000 cells/chip) and allowed to adhere O/N.

### Generation and cell transduction of lentiviral vectors

Lentiviral vectors were generated as previously described(Terrigno et al., 2018). Briefly, lentiviral vectors were prepared by transfecting HEK293T cells O/N with Lipofectamine 2000 (Invitrogen, 11668019) and a DNA mixture according to the manufacturer’s protocols. The DNA mixture consisted of the lentiviral vector of interest together with the psPAX2 packaging (Addgene #12260), the pCMV-VSV-G envelope (Addgene #8454), and the pCMV-Rev(NL4.3) (Addgene #115776) expressing plasmids in a 4:3:1:1 ratio. Transfection medium was discarded the next morning, and viral particles were collected 48 hours later and used immediately or frozen at -80°C. To achieve a high rate of cell transduction, lentiviral vectors were used fresh in a 1:1 ratio with the culture medium along with 8μg/mL Polybrene (Sigma-Aldrich, TR-1003), and cells were typically transduced O/N after passaging to increase lentiviral access to the cell surface. The lentiviral vectors used in this work consisted of the pWPXLd lentiviral backbone (Addgene #12258) containing the EGFP coding sequence. The clone the EGFP reporter with PuroR sequence, the EGFP enhancer construct was amplified by PCR using a forward and reverse primer carrying a MluI and EcoRI restriction site (forward: actacgggatccaggcctaagcttACGCGT; reverse: tagctagctactaGAATTCgagatctgagt). The vector carrying EGFP was constructed replacing the original EF1α promoter and PuroR sequence in the pLV-EF1ⲁ-IRES-Puro vector (Addgene #85132) with the amplicon carrying the EGFP reporter using the MluI/EcoRI restriction sites. The ligated vector was then sequenced to ensure correct cloning of the reporter.

### Immunofluorescence (IF) analysis

For the experiment that aimed to observe the development and survival of neurons in pure or mixed cultures, Cyclopamine and SAG-treated cells were transduced with the lentivirus carrying EGFP reporter at DIV7 and selected through the PuroR resistance.

At DIV17, both EGFP-labeled and not-labeled neurons were detached and mixed together in order to have the two pure conditions (only Cyclopamine- and only SAG-treated cultures) and the other two ones (80.20 and 50.50), together with only 1% of EGFP-positive cells. These cells were then seeded on PLO/msLam-treated glass (150000 cells/cm2).

These cultures were fixed with 2% PFA at two time points, at DIV21 and at DIV30 and analyzed through immunofluorescence.

PFA solution was added to the culture wells, incubated at RT for 9 minutes, followed by aspiration and 3 washes with 1xPBS at room temperature (RT), 10’ each. Cells were then permeabilized and blocked in 3%FCS, 3%BSA (Blocking Buffer solution) + 0.3% Triton at RT for 1 hour. The permeabilization buffer was then aspirated and replaced with a primary antibody solution in Blocking containing 0,1% Triton and the corresponding dilution in the table below. Primary solution was incubated at 4°C overnight. The next day, the primary antibody solution was removed, and cells were washed 3 times with 1XPBS at RT.

The secondary antibody solution was supplemented with corresponding anti-(host) secondary antibodies in the Blocking Buffer solution, diluted at 1:1000, and cells were incubated for 1 hour at RT. Secondary antibody solution was removed, and cells were washed 3 times with 1XPBS at RT. DAPI was added to the last 10’ PBS wash, diluted 1:10.000. After the final PBS wash, all PBS was aspirated, and cells were mounted in Aqua/Poly-mount (Polysciences, 18606-100) and allowed to cure before confocal acquisition.

Images were produced on a Leica Stellaris 5 or Zeiss LSM 900 confocal microscope, by acquiring z-stack images 10-15 optical slices thick, each slice 1m in thickness at 40x and 63x.

Regarding the analysis of the neurite length and the number of nodes, images were acquired with an HC PL APO 63x/1,40 OIL CS2 objective lens and they were reconstructed using FIJI/ImageJ and analyzed using specific plugins such as SNT (Arshadi et al., 2021). For the synaptic density analysis, the percentage of dendritic spines was analyzed by counting the number of spines and then normalized to the total length of fibers previously measured through SNT plugin (see Figure S3). Quantification analyses for density of glutamatergic and GABAergic vesicles were done by using the ImageJ Synapse Counter plugin (https://github.com/SynPuCo/SynapseCounter): the colocalization of vGlut1, vGlut2 and vGat puncta with Tubβ3 fluorescent signal was divided by the total area of fiber staining (μm^2^) present in that field to obtain a density measure of synaptic puncta per fiber (see Figure 2). Moreover, the counting of the percentage of vGlut1 and vGlut2 positive cells was calculated by dividing the percentage of nuclei surrounded by synaptic vesicles by the number of neurons (Tubβ3 positive cells) (see Figure S2) (Kempf et al., 2021).

Regarding the analysis of cell clusters in CYC and SAG cultures at DIV30 and DIV50 (Figure S4), we counted the number of DAPI positive cells using ImageJ’s “Analyze Particles” feature for each image (confocal images taken with a 10x objective); then, we reported the Z-stack measurements of each image.

Quantification was performed on three independent experiments and on selected fields for each sample. Statistical significance was assessed using one-way ANOVA and Student’s t-test followed by Tukey’s multiple comparison test after testing for normality and lognormality. For comparisons of the two time points, statistical significance was assessed using two-way ANOVA with Šídák’s multiple comparisons test.

**Table 5:** primary,secondary antibodies and chemicals used in this section:

| Antibody | Dilution | Host | Company | Catalog no |
| --- | --- | --- | --- | --- |
| GFP | 1:3000 | Chicken | Aves | GFP-1020 |
| mCHERRY | 1:1000 | Rat | Invitrogen | M-11217 |
| Tubulinβ3 | 1:10000 | Mouse | BioLegend | 801202 |
| PARVALBUMIN | 1:1000 | Guinea pig | Synaptic Systems | AB-2156476 |
| vGLUT2 | 1:1000 | Rabbit | Synaptic Systems | 135403 |
| vGLUT1 | 1:1000 | Rabbit | Invitrogen | 482400 |
| MAP2 | 1:6000 | Chicken | Novus | NB300-213 |
| vGAT | 1:500 | Guinea pig | Synaptic Systems | 131011C2 |

| Antibody | Fluorophore | Company | Catalog no |
| --- | --- | --- | --- |
| DONKEY Anti-Mouse IgG | Alexa fluor 546 | Invitrogen | A10036 |
| GOAT Anti-Rabbit IgG | Alexa fluor 488 | Abcam | AB150077 |
| GOAT Anti-Chicken IgG | Alexa fluor 488 | Invitrogen | A11039 |
| GOAT Anti-Guinea Pig IgG | Alexa fluor 488 | Invitrogen | A11073 |
| GOAT Anti-Rabbit IgG | Alexa fluor 546 | Invitrogen | A11035 |

| Chemicals | Company | Catalog no |
| --- | --- | --- |
| biotinylated Wisteria Floribunda Lectin (WFA) | Vector Laboratories | B-1355-2 |
| Streptavidin, Alexa Fluor™ 488 conjugate | Thermo Fisher | S11223 |

### RNA extraction and qRT-PCR analysis

Samples for RNA extraction were harvested following the same protocol used for splitting. After the centrifugation step, the supernatant was removed, and the cell pellet was processed using the NucleoSpin^®^ RNA kit (Machery-Nagel, 740955.250). RNA concentration was measured with the NanoDropTM Lite Spectrophotometer. For each RNA sample, approximately 200 ng of RNA were reverse transcribed into cDNA for qRT-PCR analysis using the Reverse Transcriptase Core Kit 300 (Eurogentec RT-RTCK-03). 8 µL of cDNA were then mixed with SensiFAST SYBR mix (12 ml, BioLine BIO-98020) and the amplification analyses were quantified with Qiagen 72-Well Rotorgene.

**Table 2.**
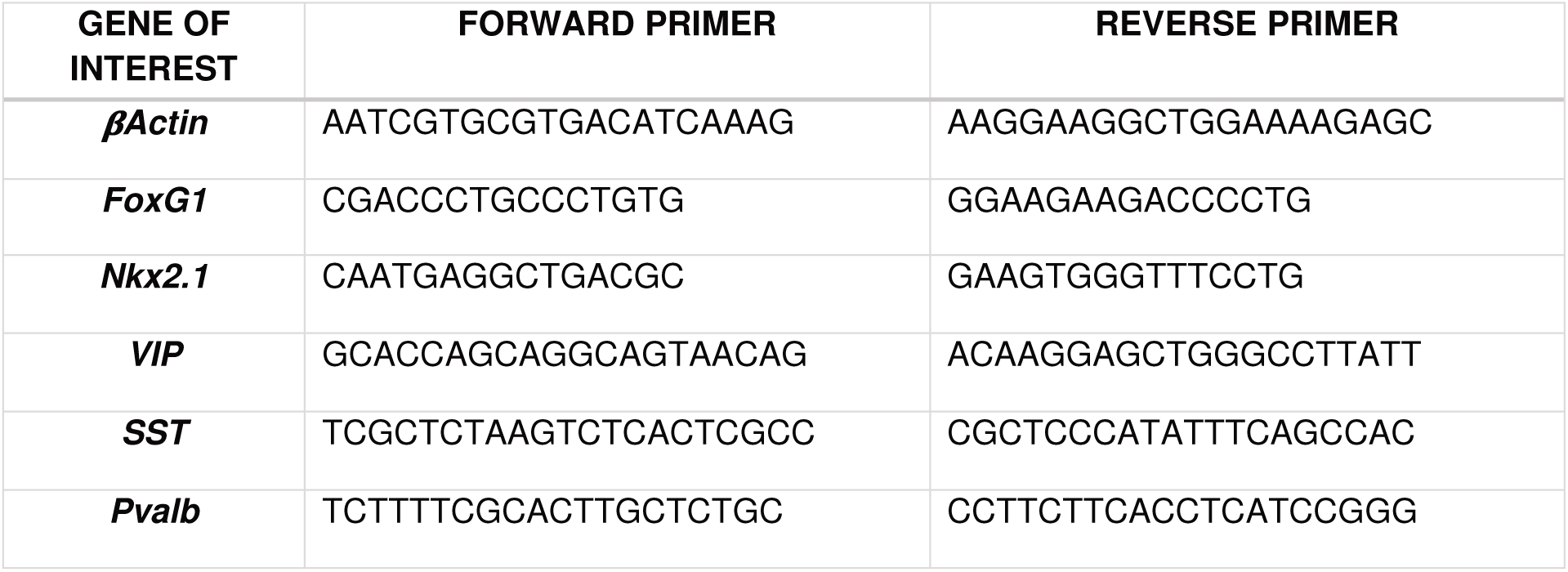
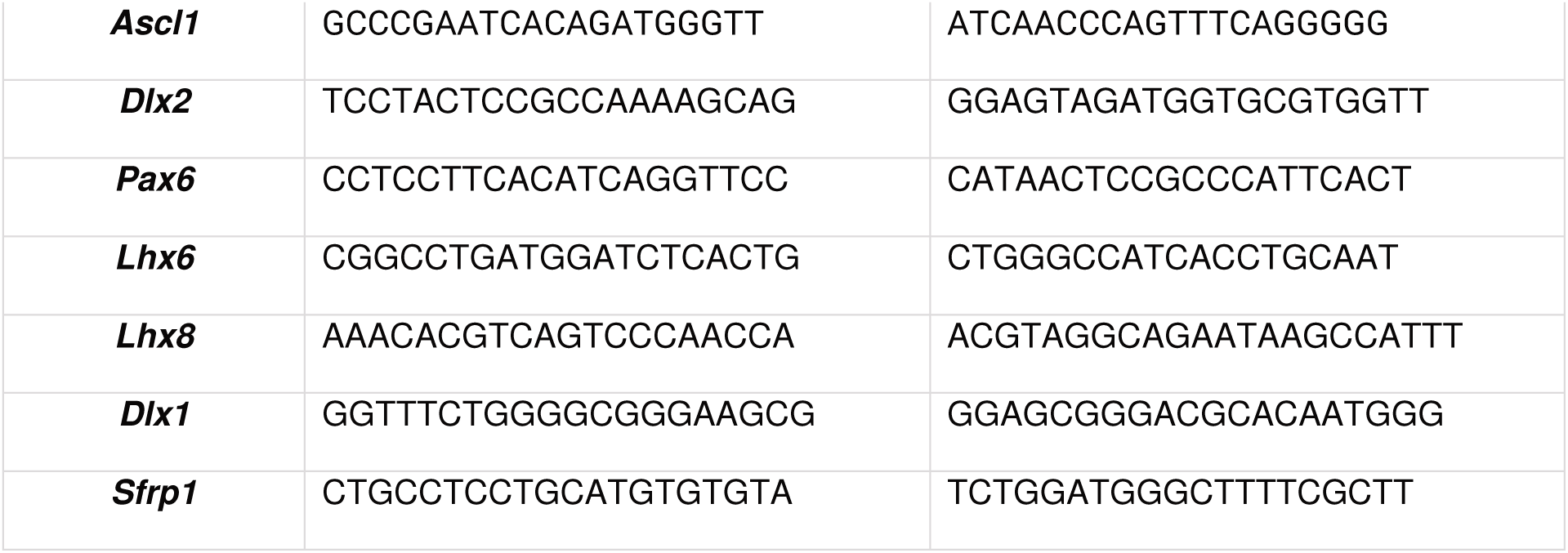
qRT-PCR primer sequences.

### qRT-PCR analysis

qRT-PCR was performed with Rotorgene devices. The method used to analyze qRT-PCR is based on the Relative Expression method supplied with the software of the Rotorgene device. The CT for each gene was obtained directly from Rotorgene. An internal control was used to reduce the variability caused by possible changes in the amount of RNA/DNA between each sample, following the ΔCT analysis method (Pfaffl, 2001). β-actin was used as the reference gene. The PCR efficiency of each sample was raised to the ΔCT to obtain the fold change of the target gene relative to the expression of β-actin (which expression was set to 1 with this method).

### Electrophysiological recordings and analysis

Neuronal cultures were seeded at DIV 18-22 onto commercially available Accura HD-MEA chips (3Brain GmbH), each equipped with 4096 CMOS microelectrodes with 60μm pitch and 21×21μm size, allowing recording of extracellular local field potentials. The 4096 electrodes are arranged in a 64×64 grid. Electrophysiological recordings were performed on DIV25-45 using the BioCam DupleX system (3Brain GmbH). After a 5 min acclimation period outside the incubator, spontaneous neuronal activity was recorded for 5 min under stable conditions (37°C, 5% CO2) and sampled at 20kHz. Spike detection was performed using BrainWave 5 software (3Brain GmbH; see next paragraph). Finally, chemical stimulation was induced by adding specific compounds to the medium: Bicuculline (BIC, 20μM; Sigma-Aldrich, 14340) to block GABA receptors, D-2-amino-5-phosphonopentanoic acid (AP5, 25μM; Sigma-Aldrich, A8054) to block NMDA receptors, and 6-cyano-7-nitroquinoxaline-2,3-dione (CNQX, 25μM; Sigma-Aldrich, C127) to block AMPA receptors. Electrophysiological activity was recorded 5 min before and 6 min after drug administration.

### Protocol of stimulation

To obtain quantitative information on the evoked network neural activity we decided to stimulate all the types of cultures. After computing the Spike Detection with PTSD algorithm, we performed an electrical stimulation on one pair of electrodes (for single stimulation) or on seven pairs of electrodes (for NBs induction) that have the highest Firing Rate (>5 spks/sec). We used a biphasic stimulus with the following parameters: current amplitude of 10 µA per electrode, with a duration of 100 µs (50 % duty cycle) and an interphase delay of 10 µs. We developed a protocol with 25 stimuli at 0,1 Hz. To evaluate network-induced activity, we performed network burst (NB) detection and classified a network burst as electrically induced if the absolute difference between its start time and the stimulus time was less than twice the time bin used in the NB detection algorithm. Then we computed the temporal duration for the spontaneous and the induced NB.

### Dispersion Index and Distribution of the global response to the stimulus

The results shown in Fig. 7 and S7 are based on the computation of temporally aggregated neural activity in response to a given stimulus.

Specifically, when a stimulation pulse was delivered at some target location of the MEA grid, electrical activity was recorded from every other MEA electrode. A spike sequence was then extracted by means of a spike detection algorithm (described in the following subsection).

Some spurious spikes and electrical artifacts where removed by means of a filtering algorithm, removing spike signals with amplitude *V ∨ _max_* above a threshold *V_thr_ = 1000mV*. Another filtering algorithm calculates the area under the spike curve *A* and divides it by the spike amplitude in order to obtain an estimate of the spike temporal width, and filters out the spike when this value is above another threshold *W_thr_ = 25us*:

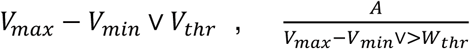

The response to a stimulus observed on a given electrode is defined as the difference in spike count between a time window immediately after the stimulus instant, and one immediately preceding it. The window size W is chosen to range from 5 to 150 ms. Mathematically, the response r_i_ over electrode i (i=1, 2, …, 4096) can be expressed as:

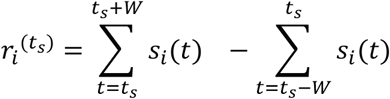

Where t_s_ denotes the stimulus delivery time, and s_i_(t) is the spike signal recorded from electrode i, i.e. a function that takes value 1 when t corresponds to the time of a spike event, and 0 everywhere else.

During a recording session, stimulations were repeated multiple times, in order to account for statistical variability. Therefore, from the recordings, multiple values of the response variable over each electrode are obtained. Calling the stimulation instants t_0_, t_1_, …, t_k_, it is possible to obtain responses r_i_^(t_0)^, r_i_^(t_1)^, …, r_i_^(t_k)^. We used k=25 in our experiments. Each of these values can be seen as a realization of a random variable R_i_, of which we wish to provide a statistically grounded estimate. This is by computing the sample mean and the corresponding confidence intervals from the observations r_i_^(t_0)^, r_i_^(t_1)^, …, r_i_^(t_k)^:

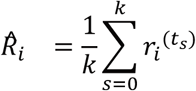

Concerning the confidence intervals, initially we employed both a parametric and a nonparametric estimator. However, we found the non-parametric estimator to be ill suited for scenarios with very few spikes in the time window, so we resorted to the parametric estimator instead. These estimator is simply based on the T-Student estimation for the confidence intervals of the sample mean:

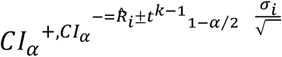

where *σ_i_* is the sample standard deviation of the observations r_i_^(t_0)^, r_i_^(t_1)^, …, r_i_^(t_k)^, and *t^k-1^_1-α⁄2_* is the required T-Student’s percentile.

Each pixel of the matrices in Fig. S7 represent the corresponding value *R̂_i_* measured for each electrode i, and represented graphically in grayscale: darker colors represent weaker responses, while brighter colors correspond to stronger responses. Moreover, when the measured confidence intervals indicate that the response value of a given pixel is significantly above zero in a statistical sense, then the corresponding pixel is colored in red. Specifically, the required confidence level is set to 95%. Similarly, when a confidence level indicates that the response of an electrode is significantly below zero, the pixel is denoted in yellow. In this case we require the positive tail of the distribution to be below zero at the 100 - 95 = 5 percentile.

Finally, the dispersion index in Fig. 7 is obtained by considering the geometric dispersion of the electrodes with significant response to a stimulus. Ideally, if all the response is concentrated in a very localized portion of the MEA grid, the corresponding dispersion should be low, while if the response is spread out all over the grid, the dispersion is large. More formally, the dispersion index is computed by considering the positions of the significantly responding electrodes over the grid, represented as x-y coordinates in the domain [0, 1]x[0, 1] (where coordinates 0, 0 denote the top-left corner, and 1-1 denote the bottom right corner of the MEA grid). Let’s denote with (x_i_, y_i_) the coordinates of electrode i, and let R = {i_1_, i_2_, …} be the set of significantly responsive electrodes. It is possible to evaluate the distance between any pair of such electrodes as

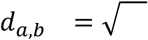

This can be again considered as a realization of a random variable D, which depends on the particular pair a, b that was chosen. It is also possible to obtain several samples of D by selecting all possible electrode pairs from R. From all these sample, we can once more evaluate the sample mean D, and the corresponding confidence intervals, for statistical comparisons.

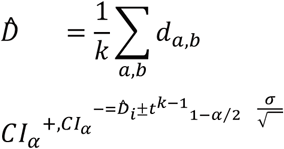

In this case, k=*R ∨ (∨ R ∨ −1)*/2, and *σ* is the sample standard deviation of the observations *d_a,b_*.

The parametric estimation methods discussed above are based on the central limit theorem, and rely on the assumption of variance finiteness in the random variables being statistically estimated. This assumption was checked by observing the homoscedasticity of the observations, i.e. sample variance converging stably as the number of observations increased.

### Spike and Burst detection

The spike detection algorithm used is the PTSD (precision time spike detection) (Maccione et al., 2009) that requires 3 parameters:

- Noise threshold (set to 10 times the standard deviation of the baseline noise for each channel individually).
- Peak lifetime period (set to 2 ms), corresponding approximately to the spike duration.
- Refractory period (set to 2 ms), which corresponds to the minimum time interval between one spike and the next (Parodi et al., 2023).

After spike detection, each channel burst was detected based on the channel spike train. The burst detection was performed as proposed in the literature (Chiappalone et al., 2005). The implemented algorithm requires 2 parameters: the maximum inter-spike interval (ISI) between two consecutive spikes of a burst (*maxiSi*, set to 50 ms) and the minimum number of spikes in a burst (*minspk*, set to 5 spikes).

Spike bursts are defined as sequences of spikes with ISI smaller than *maxiSi* and containing at least a number of spikes equal to *minspk*. The values of 50 ms and 5 spikes for these two parameters were set after a series of comparisons between the results of the burst detection algorithm and visual inspection of various experimental recordings. Mean firing rate (MFR) and mean burst rate (MBR) were calculated by counting the average number of spikes or bursts in 5 minute recordings divided by the number of active channels (firing rate greater than 0.1 spikes/s for spiking activity and burst rate greater than 0.3 bursts/min for bursting activity). To fully characterize the bursting activity, the mean burst duration (MBD), the percentage of bursting electrodes and the percentage of random spikes were calculated. The mean burst duration (MBD) was calculated as the average temporal length of the bursts. The percentage of bursting electrodes was determined as the proportion of active channels exhibiting bursting activity. Lastly, the percentage of random spikes was computed by considering all spikes that were not part of any burst activity. All these quantities were calculated for all replicates of the same culture and the Mean ± SEM was plotted over time (Figure 4).

### Network Burst activity

Network bursts (NBs) are events of collective synchronization within the culture. To quantify the level of synchronization of the neuronal network activity, we derived the mean network burst rate (NBR, number of network events per minute) and the network burst duration (NBD) for each culture. A NB was identified when the activity is composed by at least 50 consecutive spikes within a 50 ms window and the firing rate (number of spikes per bin) exceeded a threshold determined based on the mean and standard deviation of the firing rate signal. NB detection was performed as follows:

- The network’s firing rate (number of spikes per bin) was calculated using a time bin of 50 ms.
- Firing rate peaks were considered if the local firing rate maxima were greater than the mean of the firing rate signal plus 4 times the standard deviation.
- The beginning and end of each NB were defined such that the firing rate before and after each peak fell below the mean of the firing rate signal plus 2 times the standard deviation (in cases where two or more adjacent local maxima correspond to the same onset, only one event is detected).

NBR was calculated for all replicates of the same culture and the Mean ± SEM was plotted over time.

### Center of Activity Trajectories (CATs)

To quantify the propagation of coordinated network activity, we performed Center of Activity Trajectory (CAT) (Chao et al., 2007) analysis, which calculates the spatial and temporal evolution of each NB event. CAT provides a sort of center of mass for spikes, where the location of the center of mass is the physical location of the channels in the MEA map, and mass is replaced by spike activity.

The algorithm to compute NB trajectories is as follows: firstly, the start and end points of a NB are detected (as explained in the previous section) to define the time window of interest. All active channel spikes are then considered to compute the neuronal activity trajectory in this time window. A time step *o_t_ = 20*ms, is fixed for the binning of neuronal firing. Then, the activity is computed by counting all spikes for each channel in each time window *[t, t + o_t_].* Finally, the activity trajectory is defined by : *CA(t) = ^∑ch^ ^Ach(t)(rowch,colch)^,* where *A_ch_(t)* represents the activity (counted spikes) of the channel *cℎ*at time *t* (in the corresponding time window *[t, t + o_t_]*)and *(row_ch_, col_ch_)* are the physical coordinates in the MEA map (row and column) of the channel *cℎ*.

For each NB, we computed the spatial and temporal evolution of its activity trajectory throughout the duration of the synchronized event. The time evolution (with a time window of 150 ms for each NB) is represented in the plots by a colored scale, while the onset of each NB is highlighted by a blue dot.

### Network connectivity analysis

To estimate the functional connectivity within the neural networks, we used a previously validated cross-correlation-based approach (Ullo et al., 2014), dealing with point processes of events (e.g., spike trains). More precisely, for each pair of active electrodes {*x*, *y*}, the cross-correlation function (cross-correlogram) between their spike trains was estimated as:

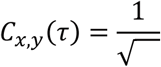

where *N_x_, N_y_* are the total number of spikes for channels *x* and *y* respectively. The time bin *L_t_* was set to 1 ms and the time delay tau (τ) was varied in a range from -2 to 2 ms. The absence of delay (τ = 0 ms) was excluded to eliminate synchronous spikes less than 1 ms apart, which are incompatible with the synaptic time delay (Poli et al., 2015). For each pair of channels, the maximum correlation value at delay τ was considered. To determine which correlations are statistically significant, it is necessary to select a threshold. We implemented a shuffling procedure that randomizes the temporal order of events to create a null hypothesis scenario where all observed correlations are purely due to chance as described in Tonelli et al., 2025.

The connectivity matrix of the randomized spike trains is computed, and the threshold is determined by computing the 99.9th percentile of the random correlation distribution. From the connectivity matrix, we derived the adjacency matrix, where the strength of the connections for the graph is represented by the correlation values (Figure 6A). Connectivity graphs were plotted while maintaining the physical position of the channels in the MEA map, with each connection colored based on the correlation value. From these graphs, we extracted the number of nodes and links (Figure 6).

### Characterization of neuronal activity after chemical drug administration

To evaluate the effect of chemical drugs on the five different cultures, we computed the probability distributions of single channel activity variation before and after drug administration. For each culture, we computed the number of spikes per channel, considering all active channels under normal and drug conditions. The drug condition was considered excluding the first 100 seconds of recording after the moment of administration. The activity variation was calculated as the differences between the channel activity after and before drug administration (number of spikes per channel), divided for the channel activity before administration. To evaluate the real effect of the drugs, for each replicate of the same culture, we calculated the variation of the baseline activity in normal condition, considering different time portions of spontaneous activity (excluding 100 seconds of recordings between one portion and another). The distributions of firing rate variation account for the activity of individual channels across all replicates. Finally, a non-parametric Mann-Whitney U test was performed to determine statistically significant differences between the baseline and post-drug distributions. Furthermore, to quantify the changes in the functional connectivity before and after drug administration, we computed the variation of the number of nodes and links compared to the baseline conditions (Figure 6C).

### Quantification and statistical data analysis

Unless otherwise stated, the data presented herein were analyzed with: GraphPad Prism software was used for statistical analysis and data plotting; Proprietary Leica and Zeiss confocal software was used for IF imaging, while Fiji software (imageJ) was used for downstream analysis; BrainWave 5 software and custom codes developed in Python were used to analyze the raw recording data of neuronal activity on the HD-MEA.

---

**Figure S1.**
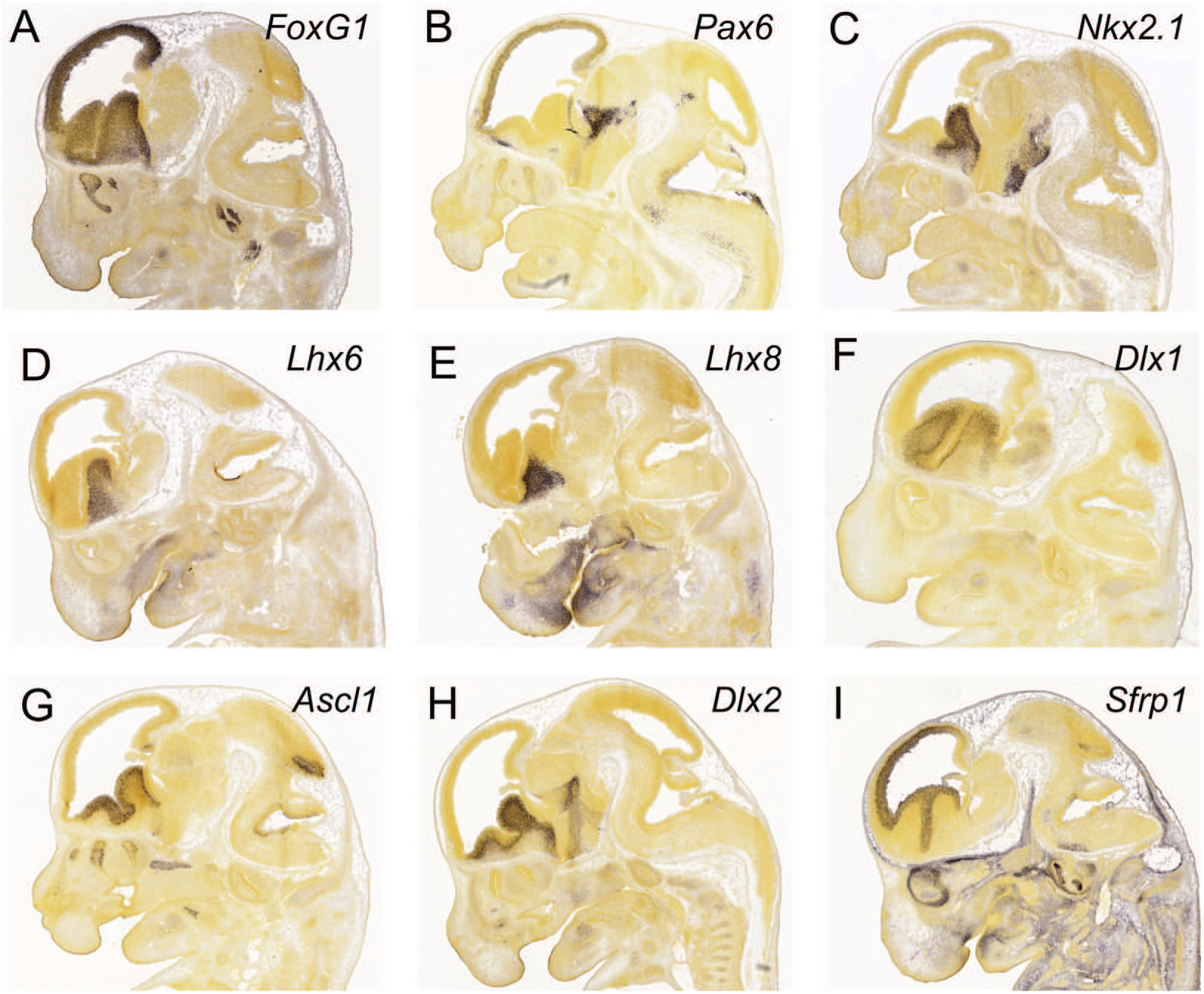
Positional markers of dorsal and ventral telencephalon. A-I) ISH of P13.5 mouse showing the expression of markers of telencephalic (A), dorsal (B), ventral (C) and subpallial (D-I) identity analyzed in Figure 1C-F. Images from Allen Brain Atlas: Developing Mouse Brain (https://developingmouse.brain-map.org/).

**Figure S2.**
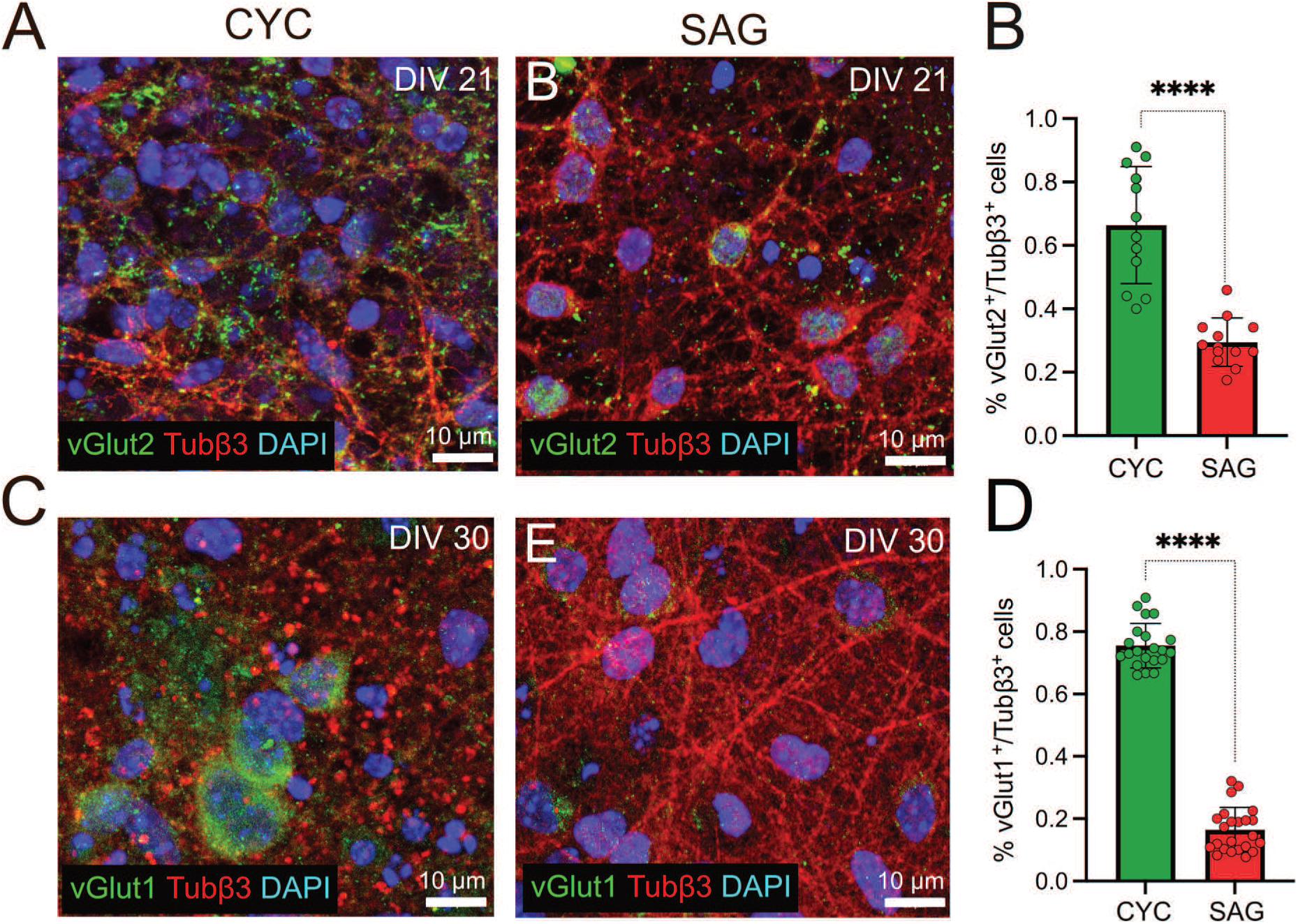
Glutamatergic marker expression in CYC and SAG cultures. A,C) Representative confocal image details of vGlut2 (A) and vGlut1 (C) positive neurons in CYC and SAG neurons at DIV21 and DIV30, stained also with Tubβ3 and DAPI. B,D) Percentages of vGlut1 (B) and vGlut2 (D) positive neurons in CYC and SAG cultures (n = 3 independent experiments), calculated by dividing the percentage of nuclei surrounded by synaptic vesicles by the number of neurons (Tubβ3 positive cells). Mean ± SD is shown, unpaired t-test, ****p-value < 0.0001.

**Figure S3.**
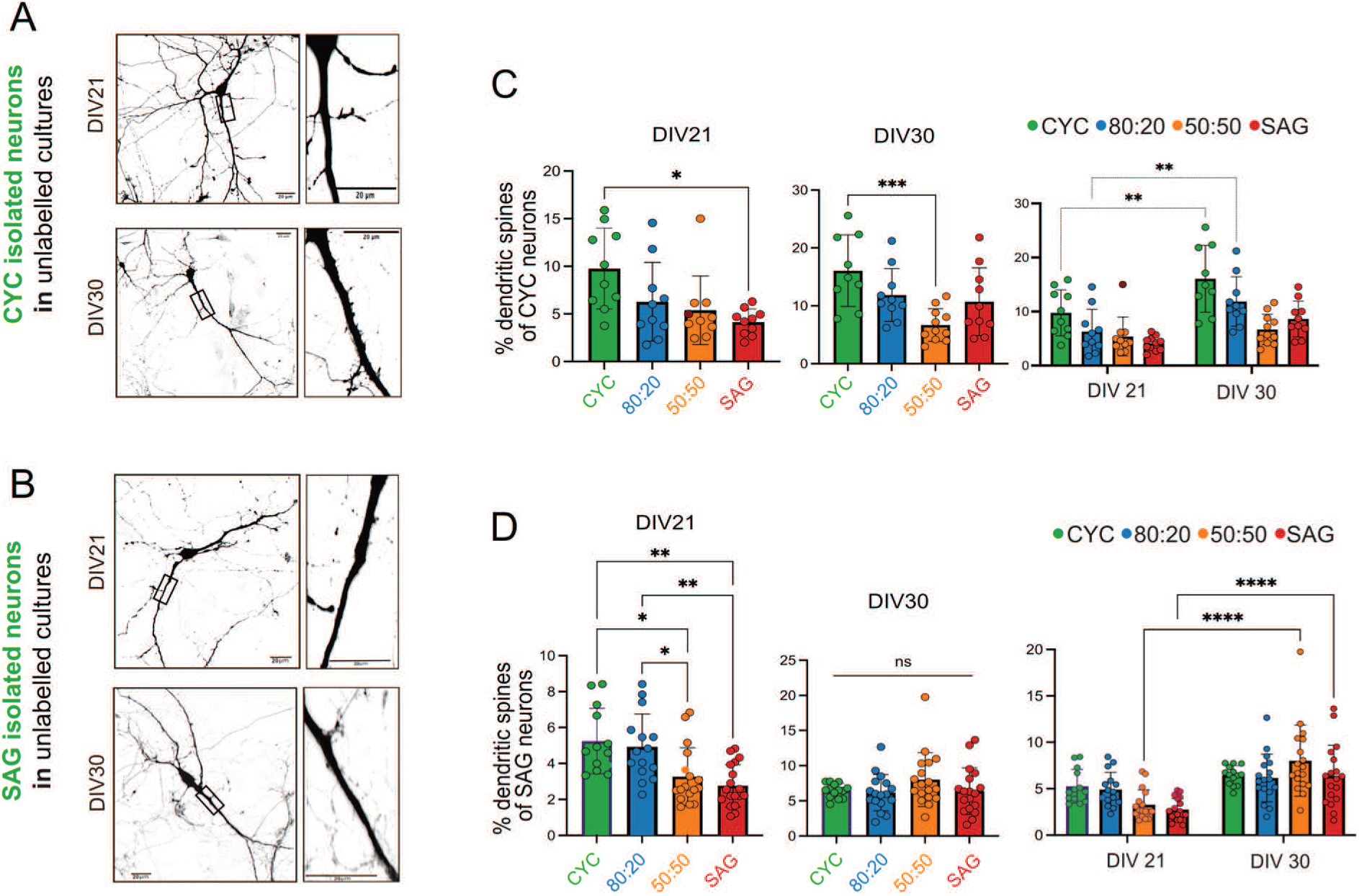
Spine analysis of CYC and SAG labeled neurons in pure and mixed CYC and SAG culture substrates. A,B) Representative confocal images of dendritic spines of CYC and SAG neurons at DIV21 and DIV30; Inset on the right: enlarged details of the main image. C,D) Quantification of the percentage of dendritic spines of CYC (C) and SAG (D) neurons at DIV21, DIV30 and over time; Mean ± SD is shown. Ordinary one-way ANOVA with Tukey’s multiple comparisons test was performed to compare samples at each time point, and two-way ANOVA with Šídák’s multiple comparisons test was performed for comparisons over time. N = 3 independent experiments; p-values: *p-value < 0.05, **p-value < 0.01, ***p-value < 0.001, ****p-value < 0.0001; ns = not significant.

**Figure S4.**
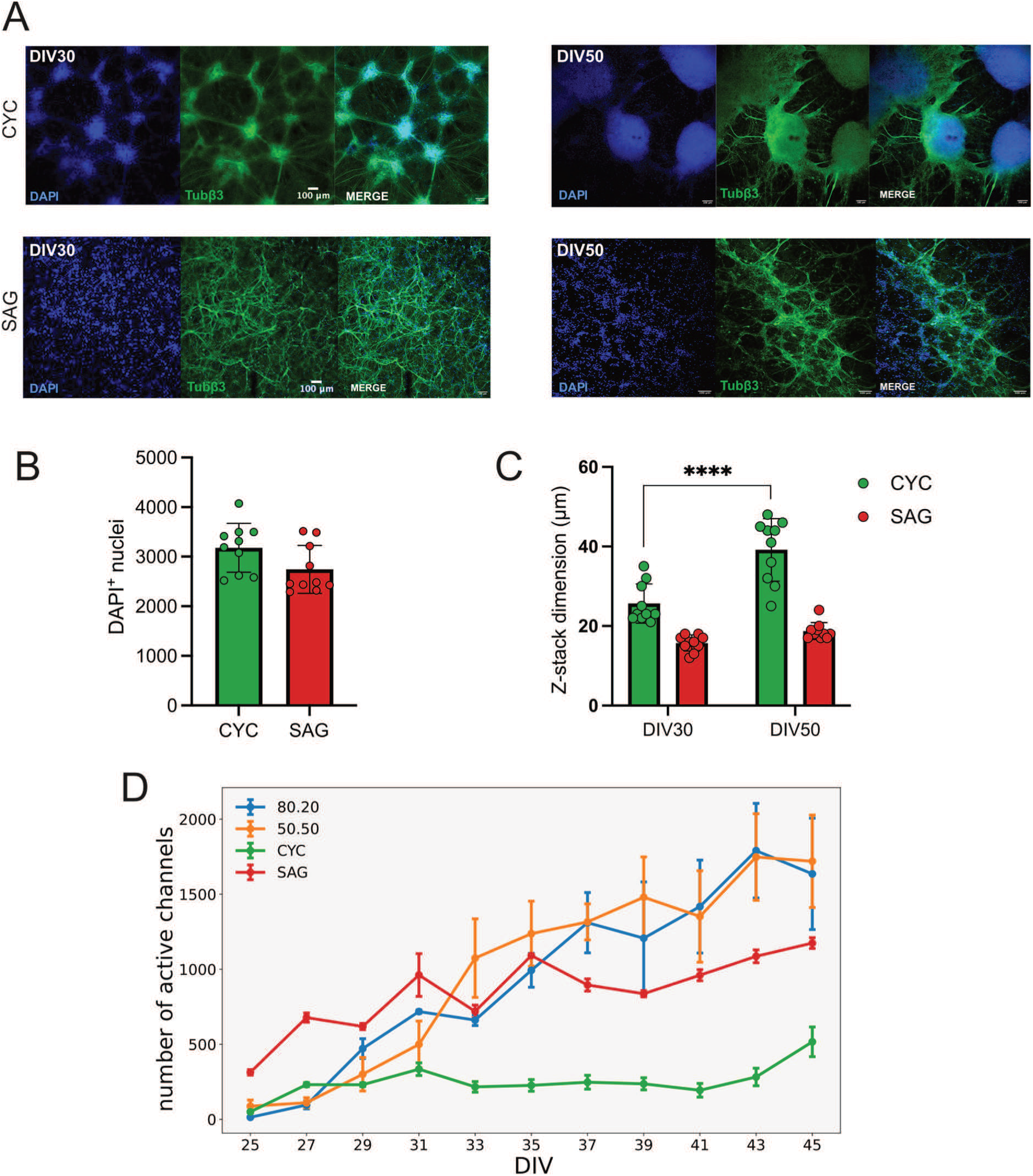
Comparison of spatial cell distribution between CYC and SAG long-term cultures. A) Representative confocal images of CYC and SAG cultures at DIV30 and at DIV50, showing morphological differences between the two cultures. B) Quantification of DAPI^+^ nuclei in each 10x image analyzed (n = 3 independent experiments); Mean ± SD is shown. C) Quantification of cluster size in CYC and SAG cultures (n = 3 independent experiments); Mean ± SD is shown; two-way ANOVA followed by Šídák’s multiple comparisons test was performed; ****p-value < 0.0001. D) Number of active channels in each condition over time (n = 3 independent experiments; Mean ± SEM is shown).

**Figure S5.**
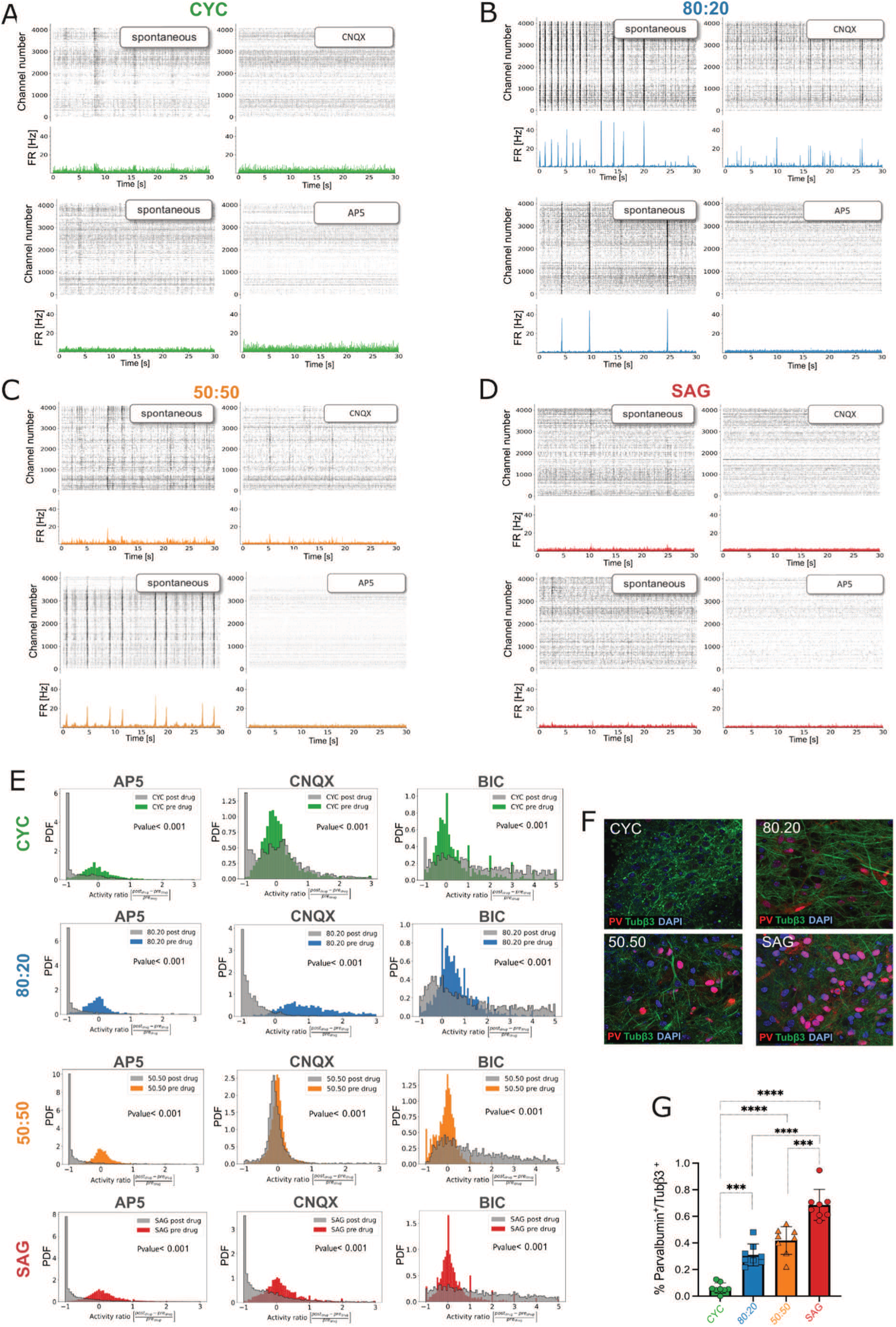
AP5, CNQX and Bicuculline significantly affect the firing activity in CYC, SAG and mixed cultures. A,B,C,D) Representative raster plots for each culture condition showing the activity before and after drug administration (AP5 and CNQX). E) Probability distribution of the activity variation of each channel, before and after drug administration. The colored distributions represent the variation of baseline activity (without drugs) considering different time portions of spontaneous activity. The gray distributions represent the variation after drug administration; Non-parametric Mann-Whitney U test between the baseline and post-drug distributions (p-values are displayed on the graph); PDF, probability density function. F) Representative confocal images of CYC, SAG, 80.20 and 50.50 cells stained with Tubβ3, Parvalbumin and DAPI at DIV35. G) Quantification of PV^+^ neurons on Tubβ3 positive cells at DIV35; Mean ± SD is shown; ordinary one-way ANOVA with Tukey’s multiple comparisons test (n = 3 independent experiments). The measured values of PV+ cells in mixed cultures (30% ± 8% in 80:20; 42% ± 10% in 50:50) are compatible with those anticipated by the mix ratios: 20% ± 9% in 80:20; 40% ± 12% in 50:50. P-values: ***p-value < 0.001, ****p-value < 0.0001.

**Figure S6.**
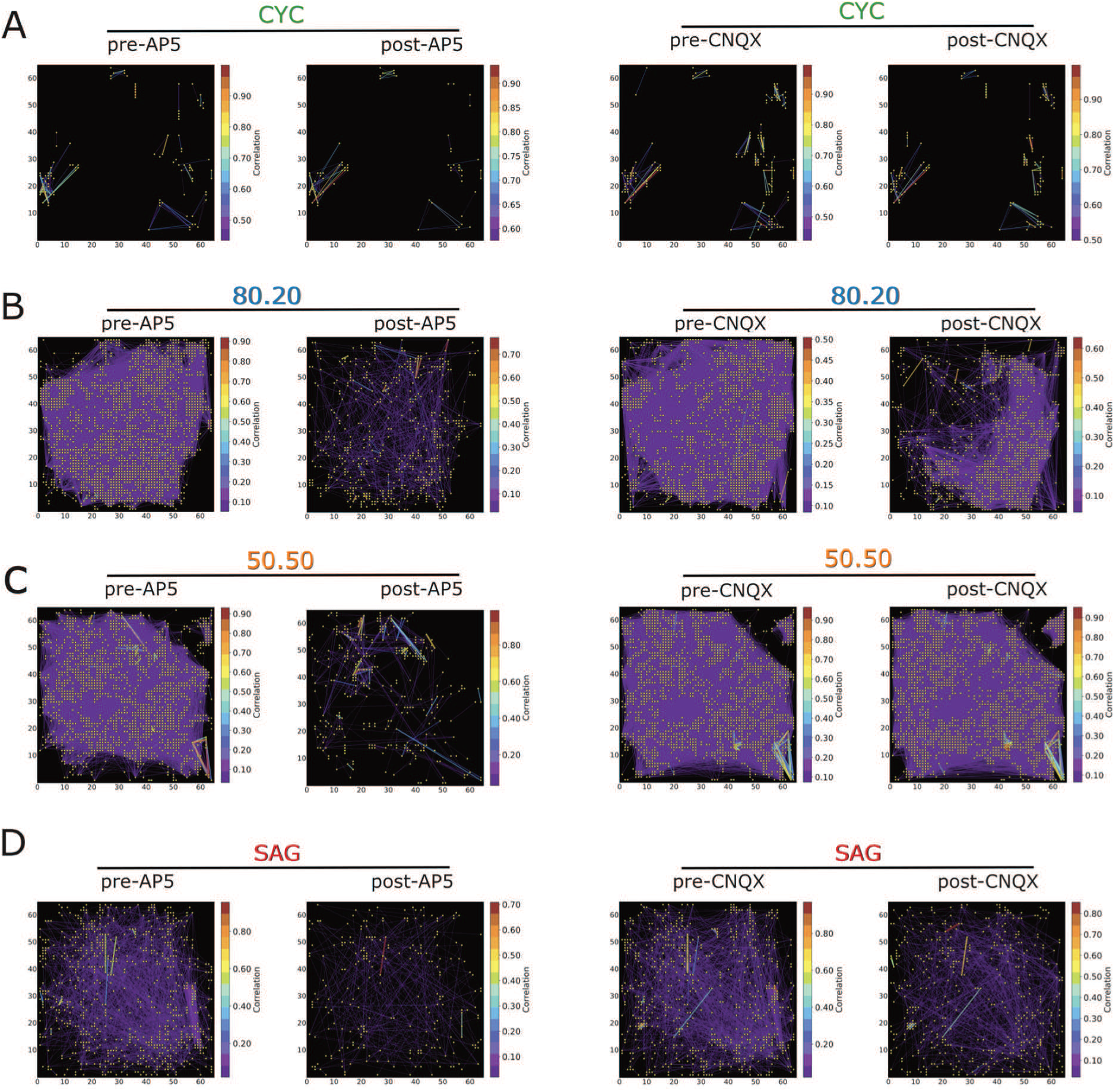
Representation of functional connectivity upon AP5 and CNQX administration. A-D) Connectivity plots of representative neuronal cultures during spontaneous activity before (pre-drug) and after (post-drug) drugs administration. Each yellow point represents a node of the functional graph; colored lines represent the correlation strength between two points (only the 10% of the functional links are shown). The color bar indicates the correlation index.

**Figure S7.**
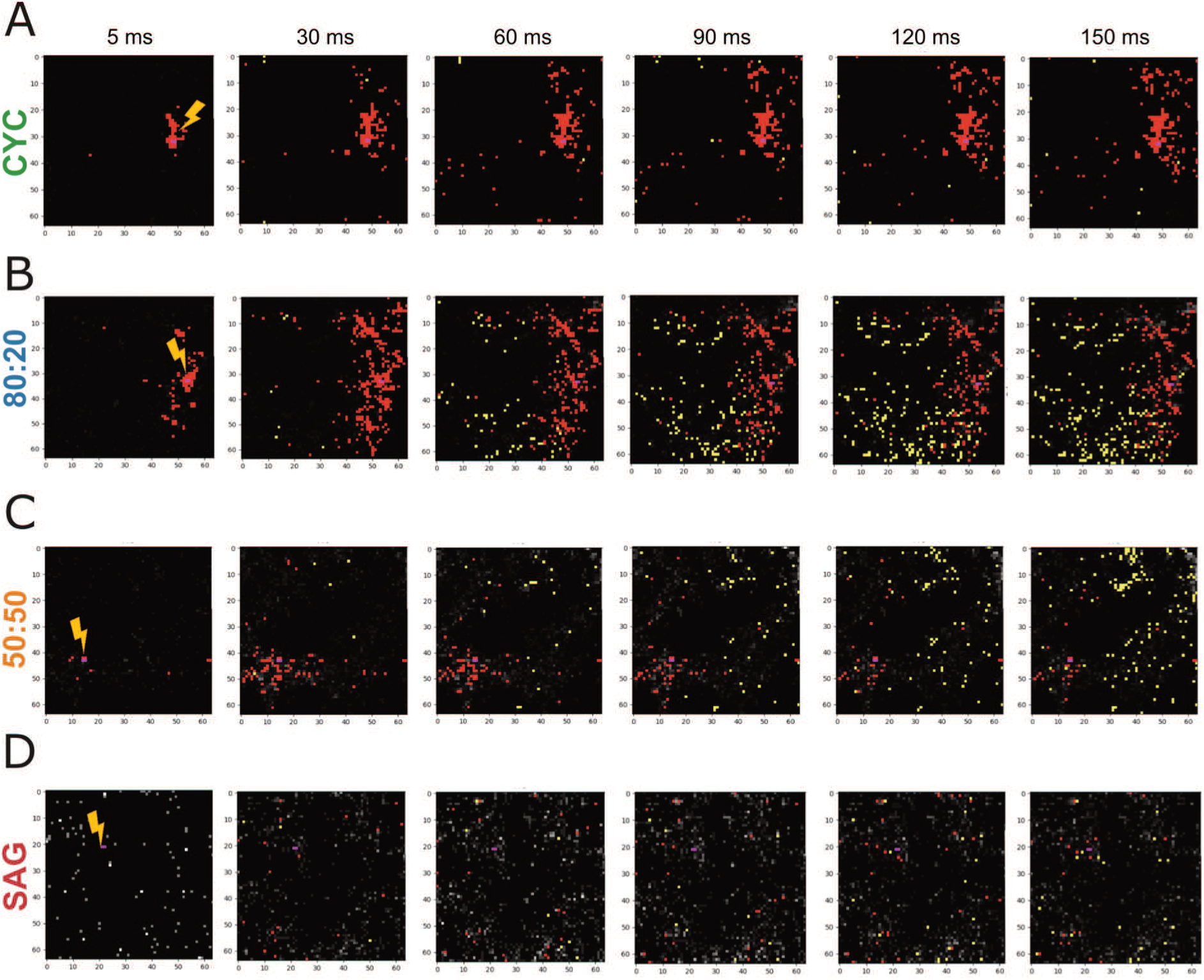
Schematic representation of single channel stimulation. A) Schematic representation of the chip plate, where each pixel represents a channel: purple pixels are two stimulated electrodes (cathode and anode), also indicated by the yellow lighting; white/grey pixels represent electrodes with baseline activity, red pixels represent channels that had a significantly increased modification as compared to the baseline activity (above the 95% Cl), while yellow pixels are those that had a significantly decreased modification as compared to the baseline activity (below 5% Cl) (See Methods).

